# Non-polio enteroviruses compromise the electrophysiology of a human iPSC-derived neural network

**DOI:** 10.1101/2025.08.27.672623

**Authors:** Feline F. W. Benavides, Syriam Sooksawasdi Na Ayudhya, Mark A. Power, Willemijn F. Rijnink, Auriane Deguergue, Bjoern Meyer, Femke M. S. de Vrij, Debby van Riel, Kristina Lanko, Lisa Bauer

## Abstract

The non-polio enteroviruses enterovirus-D68 (EV-D68) and enterovirus-A71 (EV-A71) are highly prevalent and considered pathogens of increasing health concern. While most enterovirus infections are mild and self-limiting, severe complications ranging from meningitis, encephalitis, to acute flaccid paralysis can occur, especially in children and immunocompromised patients. Despite the global burden of neurological complications caused by EV-D68 and EV-A71, the underlying neuropathogenesis remains poorly understood. In particular, the impact of the infection on neural function has not been clearly elucidated. Here, we compare the replication kinetics, cellular tropism, and electrophysiological effects of EV-D68 and EV-A71 infection in a physiologically relevant human pluripotent stem cell-derived neural co-culture model, consisting of excitatory neurons and astrocytes. Inoculation with contemporary circulating EV-D68 strains and an EV-A71 strain resulted in decreased neural activity in the co-cultures, with EV-D68 A2/2018 inducing the most rapid and robust negative effect on neural co-cultures, followed by EV-D68 B3/2019. EV-D68 strains preferentially infected neurons, whereas EV-A71 infection was detected in both cell types to the same extent. Despite the lack of viral release of infectious virus particles of EV-D68 B3/2019 in the supernatant, the infection could spread in the cultures and reduce neurotransmission. Higher viral load and broader tropism of EV-A71 did not result in enhanced impairment of neural function. Our results demonstrate that neurotropic non-polio enteroviruses lead to disruption of spontaneous neural activity in a virus-specific manner, which does not correlate with their replication efficiency.

## Introduction

The genus *Enterovirus* within the family *Picornaviridae* comprises over 250 clinically relevant virus serotypes that circulate worldwide.^1–4^ Enterovirus infections are often unnoticed and self-resolving; however, they can cause serious illnesses that are associated with major, sometimes life-threatening complications, especially in infants, young children, and immunocompromised individuals.^5–8^ Enterovirus infections display a diverse clinical spectrum, including upper and lower respiratory disease, gastroenteritis, hand-foot-and-mouth disease, conjunctivitis, myocarditis, severe neonatal sepsis-like diseases, aseptic meningitis, encephalitis, acute flaccid paralysis (AFP), and acute flaccid myelitis (AFM).^9,10^ In recent years, non-polio enteroviruses, in particular enterovirus-D68 (EV-D68) and EV-A71, have been considered pathogens of increasing health concern since their (re)-emergence is associated with the development of neurological complications.^11–13^ EV-D68 and EV-A71 are classified into several major genotypes and globally multiple EV-A71 genotypes are circulating,^14^ while among EV-D68 viruses, the subclades A2 and B3 are currently most prevalent.^15–17^ The spectrum of neurological complications for these two pathogens largely coincides and includes aseptic meningitis, encephalitis, and AFM with some differences in clinical representations.^8,10,18^

After replication in the primary infection sites – the gastrointestinal tract for EV-A71 and the respiratory tract for EV-D68 – the viruses can get access to the CNS through multiple but not mutually exclusive pathways (e.g. hematogenous spread or infection of peripheral/cranial neurons). *In vitro* studies suggest that EV-D68 and EV-A71 disseminate to the CNS via the bloodstream by disrupting the blood–brain barrier through for example direct infection of brain endothelial cells.^19^ Another option is the so-called Trojan horse mechanism, where viruses exploit blood cells to cross the blood–brain barrier. It has been shown that EV-A71 uses this mechanism; it is unclear whether EV-D68 can invade the CNS in the same way, but the infection of blood-derived immune cells has been demonstrated *in vitro.*^20,21^ In the brain, MRI studies revealed hyperintensity lesions induced by EV-D68 and EV-A71 infection, and immune responses have been detected in different brain regions, as well as the spinal cord in patients with CNS involvement.^22,23^ EV-A71 antigen was detected in neuronal cell bodies and astrocytes across different regions,^23–25^ but for EV-D68, limited pathological data are available. In the spinal cord, viral antigen and RNA of EV-D68, and EV-A71 antigen have been detected in motor neurons of the anterior horn.^24,26,27^ Among other symptoms, patients with enterovirus-induced CNS complications present with lethargy, cognitive impairment, or seizures, indicating disruption of normal neural homeostasis and function.^28,29^

Neurotransmission plays a critical role in cognitive functions, and disruptions can lead to deficits, including impairments in learning, memory, and executive function. It has been shown that viral infection can result in changes in neurotransmission, leading to changes in behavior.^30–33^ Electrophysiology is a widely used tool to investigate neurotransmission and can be used both *in vitro* and *in vivo*. A micro-electrode array (MEA) platform is a useful and easy tool to investigate the functional consequences of viral infections on neurotransmission in a continuous manner. Recently, it has been shown that EV-D68 infection of primary rat cortical neurons resulted in reduced neurotransmission measured by MEA,^34^ but research in human-derived models is lacking. Whether EV-A71 also impairs neurotransmission is currently not understood. Induced excitatory neurons generated through Neurogenin-2 (Ngn2) overexpression provide a fast and reliable model system for studying neural function.^35–37^ These Ngn2-induced neurons, when co-cultured with astrocytes, are capable of forming functional synapses and display robust electrophysiological properties. This enables us to study spontaneous neuronal activity and assess functional network connectivity in a physiologically relevant human model.^35,37–39^

In this study, we investigated the functional consequences of infection with clinically prevalent enteroviruses in a human pluripotent stem cell (hPSC)-derived neural co-culture model. To this end, we employed networks of neural co-cultures to characterize neurotropism, replication kinetics and neural activity upon enterovirus-D68 and enterovirus-A71 infection on a MEA platform to better understand the underlying mechanisms of neurological complications associated with non-polio enterovirus infections.

## Material and Methods

### Cells and Reagents

HeLa-Rh cells (kindly provided by Johan Neyts from the Rega Institute at the KU Leuven) and HEK293T cells (ATCC) were maintained in Dulbecco’s Modified Eagle’s medium (DMEM; Capricorn Scientific, Frankfurt am Main, Germany) supplemented with 10% dialysed fetal bovine serum (FBS, Sigma-Aldrich, St. Louis, MO, USA), 2mM L-glutamine (Capricorn Scientific) and 1% penicillin/streptomycin (Capricorn Scientific) at 37°C with 5% CO_2_. Medium was refreshed every 2–4 days, and cells were passaged at >80% confluence using PBS and trypsin-EDTA (Capricorn Scientific). Cells were regularly checked for presence of mycoplasma. The compounds (*S*)-fluoxetine (Sigma Aldrich), forskolin (Tocris) and rolipram (Tocris) were dissolved in DMSO at 10 mM, 25 mM and 50 mM stock concentration, respectively.

### Viruses

EV-D68 strains included in this study were isolated from clinical specimens at the National Institute of Public Health and the Environment (RIVM), Bilthoven, The Netherlands. The clinical specimens were isolated from respiratory samples and propagated on RD cells at 33°C in 5% CO2. EV-D68 viruses from different clades were included in this study, virus reference number and accession number are as follow: clade A/2012 (4311200821; Accession Number: MN954536) A2/2018 (4311400720; Accession Number: MN954537), B1/2013 (4311300117; Accession number MN954538), B2/039 (4311201039; Accession Number: MN954539), B3/2019 (3101900710; Accession Number: MN726799) and Fermon. All virus stocks were Sanger sequenced. Recombinant viruses from EV-A71 Sep006 and EV-D68 Fermon were obtained by transfecting the plasmids of the infectious clones pCAGGS-EVA71-Sep006 and pCAGGS_EV-D68-Fermon in HEK293T cells. After obtaining full cytopathic effect in HEK293T cells, the virus was one more time passaged in Hela-Rh cells.

### Human induced pluripotent stem cells

Human pluripotent stem cells (hPSC) WTC-11 (Coriell no. GM25256, obtained from the Gladstone Institute, San Francisco, CA, USA), were used to generate astrocytes. WTC-11 hPSCs were maintained in hPSCs medium (**Table 1**), released with Accutase (Life Technologies), and grown in Matrigel (Corning)-coated 6-well plates. Medium was refreshed every other day, and cells were cultured at 37°C and 5% CO2.

**Table 1.**
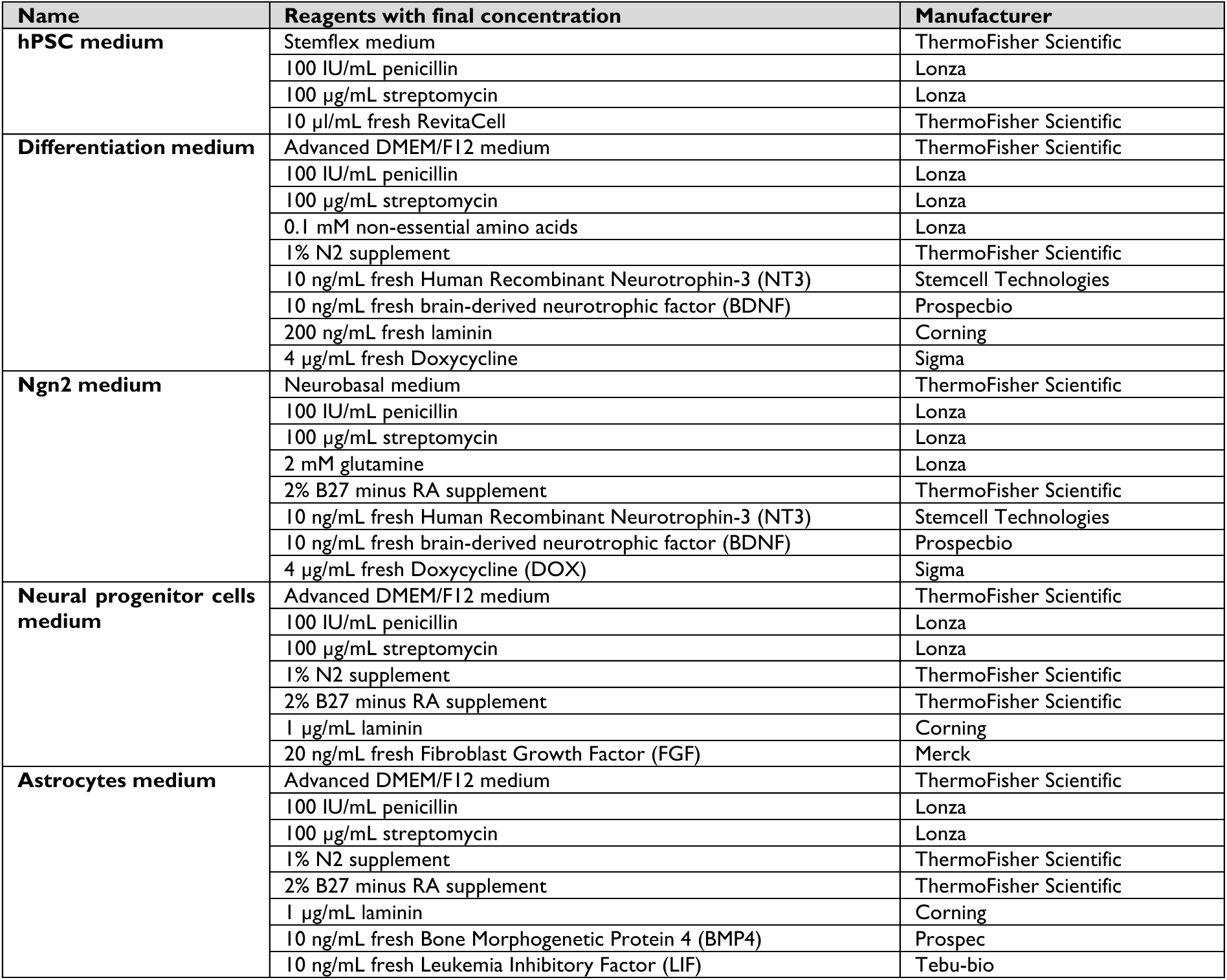
Overview of media used for differentiation and maintaining of stem cells.

### Differentiation of hiPSC into NGN2 neuron-astrocyte co-cultures

WTC-11 hPSCs were directly differentiated into excitatory cortical layer 2/3 neurons by overexpression of neurogenin-2 (Ngn2) using an adapted protocol for inducible overexpression in the presence of doxycycline, as previously described.^35,36,40^ In short, coverslips or MEA plates were coated with poly-L-ornithine (Sigma, 100 µg/mL) for 1hr at RT in the dark. Then coverslips or MEA plates were washed three times with water and dried. Coverslips or MEA plates were then coated with Matrigel (Corning, 10 µl/mL resuspended in KO DMEM) for 1h at 37 °C. After incubation, the hPSCs were plated in hPSC medium (**Table 1**) supplemented with 4 µg/mL fresh doxycycline (Sigma). The next day, the medium was refreshed with differentiation medium (**Table 1**). In order to guarantee the formation of functional synapses and thus functional synaptic plasticity within the network hPSC-derived astrocytes were added on day 3. hPSC-derived astrocytes were differentiated from hPSCs through neural progenitor cells (NPCs) as previously described.^37^ hPSC-derived astrocytes were added to the culture in a 1:1 ratio. The medium was refreshed the day after with Ngn2 medium (**Table 1**). Every other day, half of the medium was refreshed until day 21. During maintenance, and differentiation of hPSC-derived astrocytes and NPC, whole medium was refreshed with astrocytes or NPC medium (**Table 1**) every other day and cells were split once per week. All cells were kept at 37 °C and 5% CO_2_.

### Virus infection

Virus infections were performed by incubating hPSC-derived neural co-cultures with virus at a multiplicity of infection (MOI) of 0.1 at 37°C in 5% CO_2_ for 1 hour for viral titration or immunofluorescent staining. Subsequently, the inoculum was removed, the cultures were washed once with PBS after which half fresh and half old medium was added to the hPSC-derived neural co-cultures. Supernatants for virus titration were collected at the indicated time points. For MEA measurements, hPSC-derived neural co-cultures were incubated with virus at MOI 1, inoculum was removed after 1 hour, washed once with PBS and old medium was added 1:1 to the neural co-cultures. Medium was refreshed half every other day.

### Virus titration

Virus titres were determined by endpoint dilution on a subconfluent layer of Hela Rh cells. Briefly, 10-fold serial dilutions of samples were titrated on Hela Rh cells and incubated at 33°C in 5% CO_2_. At day 5, virus titres were determined by visual inspection of cytopathic effect. Viral titres were calculated according to the method of Reed and Munch and expressed as 50% tissue culture infective dose (TCID50).^41^

### Immunofluorescence staining

At indicated time points, hPSC-derived neural co-cultures were fixed with 10% formalin for 15 minutes and permeabilized with 1% triton (Sigma; T8787) in PBS for 15 minutes. Cells were blocked with 5% bovine serum albumin (BSA; Aurion) for 30 minutes after which cells were incubated with primary antibodies for 1 hr. Cells were washed twice with washing buffer (PBS with 0.1% BSA) and incubated with secondary antibodies for 1 hr at room temperature. Primary antibodies and secondary antibodies with used concentrations can be found in **Table 2**. Nuclei were stained using the dye Hoechst (1:1000, Invitrogen, H3570) for 10 minutes at room temperature. Samples were processed using a Zeiss LSM 700 laser scanning microscope. All images were processed using Zen 2010 software and Image J.

**Table 2.** Concentration of antibodies.

| **Antibody** | **Antibody** | **Concentration** | **Manufacturer** | **Catalogue no.** |
| --- | --- | --- | --- | --- |
| Primary | Anti-dsRNA Antibody, clone rj2 | 1:400 dilution | Jena Bioscience | RNT-SCI-10010200 |
|  | Rabbit anti-EV-D68 VP1 | 10 ug/mL | GeneTex | 132313 |
|  | Guinea pig anti-MAP2 | 2% v/v | Synaptic Systems | 188004 |
|  | Mouse anti-GFAP | 2.5 ug/mL | BD Pharma | 556327 |
|  | Chicken anti-GFAP | 10 ug/mL | Abcam | AB-4674 |
| | Mouse α-Homer1 | 13.33 μg/mL | Synaptic systems | 160011 |
| | Rabbit α-Synapsin I | 10 μg/mL | Synaptic systems | 106103 |
| | Rabbit Cleaved Caspase-3 | 0.5 μg/mL | Cell Signalling | 9661 |
| Secondary | Donkey anti-rabbit AF488 | 20 ug/mL | Invitrogen | A21206 |
|  | Donkey anti-mouse AF555 | 10 ug/mL | Invitrogen | A31570 |
|  | Donkey anti-guinea pig AF647 | 2.5 ug/mL | Jackson Immuno research | 706-605-148 |
|  | Donkey anti-chicken AF555 | 2 ug/mL | Invitrogen | A78949 |

### MEA recordings of neural activity in Ngn2 neurons co-cultured with astrocytes

Neural co-cultures were plated on 24-well CytoView MEA plates (Axion Biosystems). Plates were acclimatized for at least 5 minutes before neural activity and viability were measured for 3 minutes using the Axion Biosystems Maestro MEA at 37°C and 5% CO_2_. Data analysis was performed using AxIs software (Axion Biosystems Inc.). Covered electrodes were defined as electrodes with a minimum resistance of 18 kΩ. We excluded wells for further analysis if a well reached ≤5 covered electrodes at any point during the experiment. Active electrodes were defined as electrodes with a minimum of five spikes per minute. We excluded wells for analysis if a well had ≤5 active electrodes at the baseline recording for further analysis. The threshold for spike detection was defined as ≥6-fold the standard deviation (SD) of the root mean square noise. The threshold for burst detection was defined as >5 spikes within a time window of 100 ms. The threshold for network burst detection was defined as >50 spikes within a time window of 100 ms, with a minimum of 50% participating electrodes per well.

For stimulation and inhibition experiments, pre-warmed drugs were infused in the medium in an amount of maximum 10% of the medium. Before adding drugs, a baseline recording was performed. For stimulation, protocol was followed as described before.^42^ In short, 50 µM forskolin and 0.1 µM rolipram were diluted in Ngn2 medium. The medium containing forskolin and rolipram was added to the cells and were incubated for half an hour, before washing it out. For inhibition, a protocol was followed as described before.^43^ 10 µM (*S*)-fluoxetine (SFX) was diluted in Ngn2 medium and added to the neural co-cultures. As negative control, medium containing the same amount of DMSO was included.

### PCA analysis and heatmap generation

MEA datasets from neural co-cultures (mock-inoculated or inoculated with EV-D68 A2/2018, EV-D68 B3/2019, or EV-A71 Sep006) were first normalized to baseline recordings obtained prior to inoculation. Only data that met the inclusion criteria described earlier for MEA recordings were used. Any experiment that contained *Not Available* values throughout any of the chosen MEA variables (**Table 3**) was excluded for PCA analysis, to ensure biological relevance and integrity of the dataset. Data were mean-centered and scaled to unit variance (z-score normalization). PCA was performed using the prcomp function in R (v4.5.1; RStudio) with the stats package, and results were visualized with ggplot2 (v3.5.2). The first two principal components (PC1 and PC2) were plotted with a 95% prediction ellipse. Heatmaps were generated using pheatmap (v1.0.12) on the z-scored data matrix. Hierarchical clustering was performed on both samples and features using Euclidean distance and complete linkage, with dendrogram-based ordering. Data processing used readxl (v1.4.3) and dplyr (v1.1.4). Color intensity reflects a positive or negative effect on neural activity, with white to red indicating a more positive effect.

**Table 3.** Output MEA variables that were chosen for Principal Component Analysis as defined by the Axion Neural Metric guide.

| MEA variables | Definition |
| --- | --- |
| Mean Firing Rate | Total number of spikes divided by the duration of the recording |
| Mean inter-spike interval (ISI)<br>Coefficient of Variation | Measure of spike regularity: the coefficient of variation (standard deviation/mean) of the inter-spike interval |
| Number of Active Electrodes | Number of electrodes with activity greater than 5 spikes per minute (inclusion criteria) |
| Number of Bursting Electrodes | Number of electrodes within the well with bursts that are included in analysis (inclusion criteria for bursts: >5 spikes within 100 ms/electrode) |
| Burst Duration | Time from the first spike to last spike in a single-electrode burst |
| Number of Spikes per Burst | Number of spikes in a single-electrode burst |
| Mean ISI within Burst | ISI time between spikes, for spikes in a single-electrode burst |
| Median ISI within Burst | Median ISI time between spikes, for spikes in a single-electrode burst |
| Inter-Burst Interval (IBI) | Time elapsed between the end of one burst and the beginning of the next |
| Burst Frequency | Total number of single-electrode bursts divided by the duration of the recording |
| IBI Coefficient of Variation | Measure of burst regularity: the coefficient of variation (standard deviation/mean) of the inter-burst interval |
| Burst Percentage | The number of spikes in single-electrode bursts divided by the total number of spikes, multiplied by 100 |
| Network Burst Frequency | Total number of network bursts divided by the duration of the recording (included when $\geq 50\%$ electrodes are participating in the burst and the burst have >50 spikes within 100 ms) |
| Network Burst Duration | Average time from the first spike to last spike in a network burst |
| Number of Spikes per Network Burst | Number of spikes in a network burst |
| Number of Electrodes Participating in Burst | Number of electrodes with activity during a network burst |
| Number of Spikes per Network Burst per Channel | Number of spikes in a network burst divided by the number of electrodes participating in that burst |
| Network Burst Percentage | The number of spikes in network bursts divided by the total number of spikes, multiplied by 100. |
| Network ISI Coefficient of Variation | Measure of spike regularity across network activity: the coefficient of variation (standard deviation/mean) for the inter-spike interval for all spikes on all electrodes in a well |
| Network IBI Coefficient of Variation | Measure of network burst regularity: the coefficient of variation (standard deviation/mean) for the inter-network burst interval, the time between network bursts. |
| Network Normalized Duration IQR | Measure of network burst duration regularity: interquartile range of network burst durations. |
| Area Under Normalized Cross-Correlation | Area under the well-wide pooled inter-electrode cross-correlation normalized to the auto-correlations. Higher areas indicate greater synchrony |
| Width at Half Height of Normalized Cross-Correlation | Measure of network synchrony: distance along the x-axis (phase lag) from left half height to right half height (probability) of the normalized cross-correlogram. |
| Synchrony Index | A unitless measure of synchrony between 0 and 1.<sup>44</sup> Values closer to 1 indicate higher synchrony. |
| Resistance - Avg (kOhms) | Resistance is a measure of viable cell coverage over the electrode. Higher values indicate more intact cells are attached to the electrode (inclusion when resistance >18 kΩ) |
| Number of Covered Electrodes | Covered electrodes are defined as electrodes with resistance greater than 18 kΩ |

### Statistical analysis

Statistical differences between experimental groups were determined as described in the figure legends. P values of ≤ 0.05 were considered significant. Graphs and statistical tests were performed with GraphPad Prism version 10.5.0. Figures were prepared with ImageJ 1.54f and Adobe Illustrator 27.8.1.

## Results

### Characterization of hPSC-derived neuron and astrocyte co-cultures to study the neurovirulence of non-polio enteroviruses

To study the effect of enterovirus infection on neural networks, we employed a rapid differentiation protocol that yields hPSC-derived excitatory glutamatergic cortical neurons by overexpressing the transcription factor Ngn2 as previously described^35,36^ (**Supplementary Fig. 1**). We supplemented the Ngn2-induced neurons with hPSC-derived astrocytes to ensure neuron survival and maturation (**Supplementary Fig. 1A**). Immunofluorescence staining revealed that the cultures contained GFAP^+^ astrocytes and MAP2^+^ neurons at 21 *days in vitro* (DIV) (**Fig. 1A**). Furthermore, immunofluorescence staining of pre- and postsynaptic density proteins showed that these cultures establish synaptic connections (**Fig. 1B**). We next assessed neural network activity of the Ngn2 neural co-cultures using a MEA system (**Fig. 1C, Supplementary Fig. 1B**). The neural co-cultures exhibit spontaneous neural activity, displayed by firing rate, burst frequency, and network burst frequency between DIV 21 and 31 (**Fig. 1D**). To probe the activity of the cultures, we stimulated the neural co-cultures with 50 µM forskolin (FSK) and 0.1 µM rolipram (ROL) to induce long-term potentiation (**Fig. 1E**). A significant decrease in firing rate and a significant increase in the burst and network burst frequency were observed upon treatment, similar to previous reports^42^ (**Fig. 1F**). In addition, we aimed to inhibit the neural activity of the Ngn2 neural co-cultures. To achieve this, we used the S-enantiomer of fluoxetine (SFX), as it has neuro-modulatory effects as well as neurotoxic effects in primary rat cortical neurons.^43^ Adding 10 µM of SFX to the neural co-cultures potently inhibited the neural network activity (**Fig. 1E and 1F**). Together, this shows that the neural co-culture model is suitable to detect changes in the network activity.

**Figure 1.**
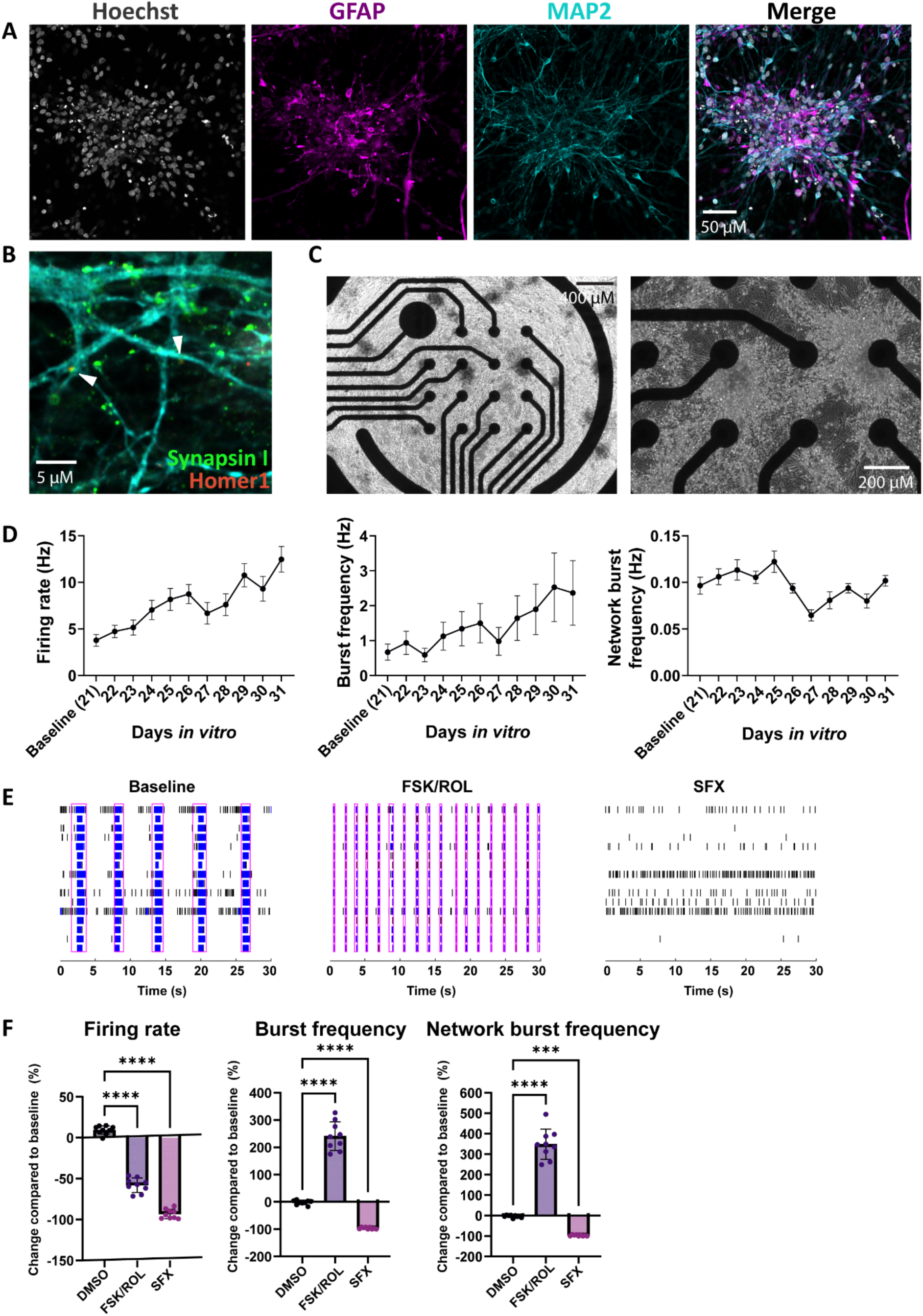
Ngn2-expressing neurons co-cultured with astrocytes form functional neural networks and respond to stimulation and inhibition. hPSC-derived Ngn2-expressing neurons were co-cultured with hPSC-derived astrocytes and matured for 21 DIV. (A) Immunofluorescent staining was performed with astrocytic-specific marker GFAP (magenta) and the neuron-specific marker MAP2 (cyan). Cells were counterstained with the nuclear dye Hoechst at DIV 21. (B) Immunofluorescent staining for functional synapses was performed against MAP2 (cyan), synapsin 1 (green, marker for pre-synaptic density), and Homer 1 (red, marker for post-synaptic density) at DIV 22. (C) Brightfield images of neural co-cultures plated on micro-electrode array plates to measure neural activity at day 21. (D) Parameters for neural activity of Ngn2 neural co-cultures between DIV 21 and 31: firing rate, burst frequency, and network burst frequency. Data are depicted as mean ± SEM and are derived from eight independent experiments (*n* = 48, unless datapoints were excluded based on exclusion criteria, see Material and Methods). (E) Raster plots of neural co-cultures were stimulated with 50 µM FSK and 0.1 µM ROL or inhibited with 10 µM SFX. (F) Firing rate, burst frequency, and network burst frequency measured after stimulation or inhibition are shown. Data are depicted as mean ± SD. Data are derived from two independent experiments (*n* = 9 per group, unless datapoints were excluded based on exclusion criteria, see Material and Methods). Abbreviations: hPSC = human induced pluripotent stem cells; Ngn2 = Neurogenin-2; DIV = days *in vitro*; GFAP = glial fibrillary acidic protein; MAP2 = microtubule-associated protein 2; SEM = standard error of the mean; DMSO = dimethylsulfoxide; FSK = forskolin; ROL = rolipram; SFX = (*S*)-fluoxetine; SD = standard deviation.

### Non-polio enteroviruses infect neurons and astrocytes and do not induce the cleavage of caspase-3

To study the neurotropism and replication kinetics, neural co-cultures were inoculated with EV-D68 strains A/2012, A2/2018, B1/2013, B3/2019, the cell-culture adapted B2/039^45^, and a recombinant Fermon virus, as well as a recombinant EV-A71 Sep006 from genotype C4 at a MOI of 0.1. At 1-, 24-, 48-, and 72-hours post inoculation (hpi), viral titres were determined in the supernatants by endpoint dilution. All viruses, except EV-D68 B3/2019, released infectious virus particles into the supernatant, suggesting efficient replication in the neural co-cultures (**Fig. 2A**). We observed that, despite comparable EV-D68 and EV-A71 inputs, EV-A71 showed a 1.5 log higher virus titre at 1 hpi after washing the cultures three times. Overall, EV-A71 Sep006 replicated more efficiently compared to all EV-D68 strains based on an area under the curve analysis after adjusting for the higher virus titer 1 hpi (**Supplementary Fig. 2**). Notably, few of the neural co-cultures began detaching from the wells between 48 and 72 hpi, with some wells completely detaching. This effect was more pronounced in EV-D68-inoculated cultures compared to EV-A71-inoculated cultures. Additionally, we observed small, darker, and denser areas in the dendrites upon infection with all viruses, likely representing protein aggregation in the dendrites of inoculated neural co-cultures, similar to what was observed in EV-D68-infected hPSC-derived motor neurons.^46^ To determine the cellular tropism of EV-D68, neural co-cultures were fixed at 24 and 72 hpi and stained for the structural protein VP1 **(Fig. 2B and Supplementary Fig. 3A**). Even though we did not detect EV-D68 B3/2019 titres in the supernatants, virus-infected MAP2^+^ cells were detected. The cell tropism of EV-A71 was visualized using double-stranded RNA, as a marker for active viral replication (**Figure 2C and Supplementary Fig. 3B**). All EV-D68 strains infected predominantly MAP2+ cells and only sparsely GFAP+ cells **(Supplementary Fig. 4A)**. EV-A71 did not show a cellular preference for MAP2^+^ neurons or GFAP^+^ astrocytes, but infected both cell types equally (**Supplementary Fig 4B**). A spotty and condensed pattern of GFAP^+^ staining was observed after inoculation with EV-D68 and EV-A71, especially after 72 hpi. Taken together, all enteroviruses showed efficient replication in Ngn2 neural co-cultures, although B3/2019 was impaired in viral release. While all EV-D68 viruses showed a clear preference for neurons, EV-A71 infected both cell types equally.

**Figure 2.**
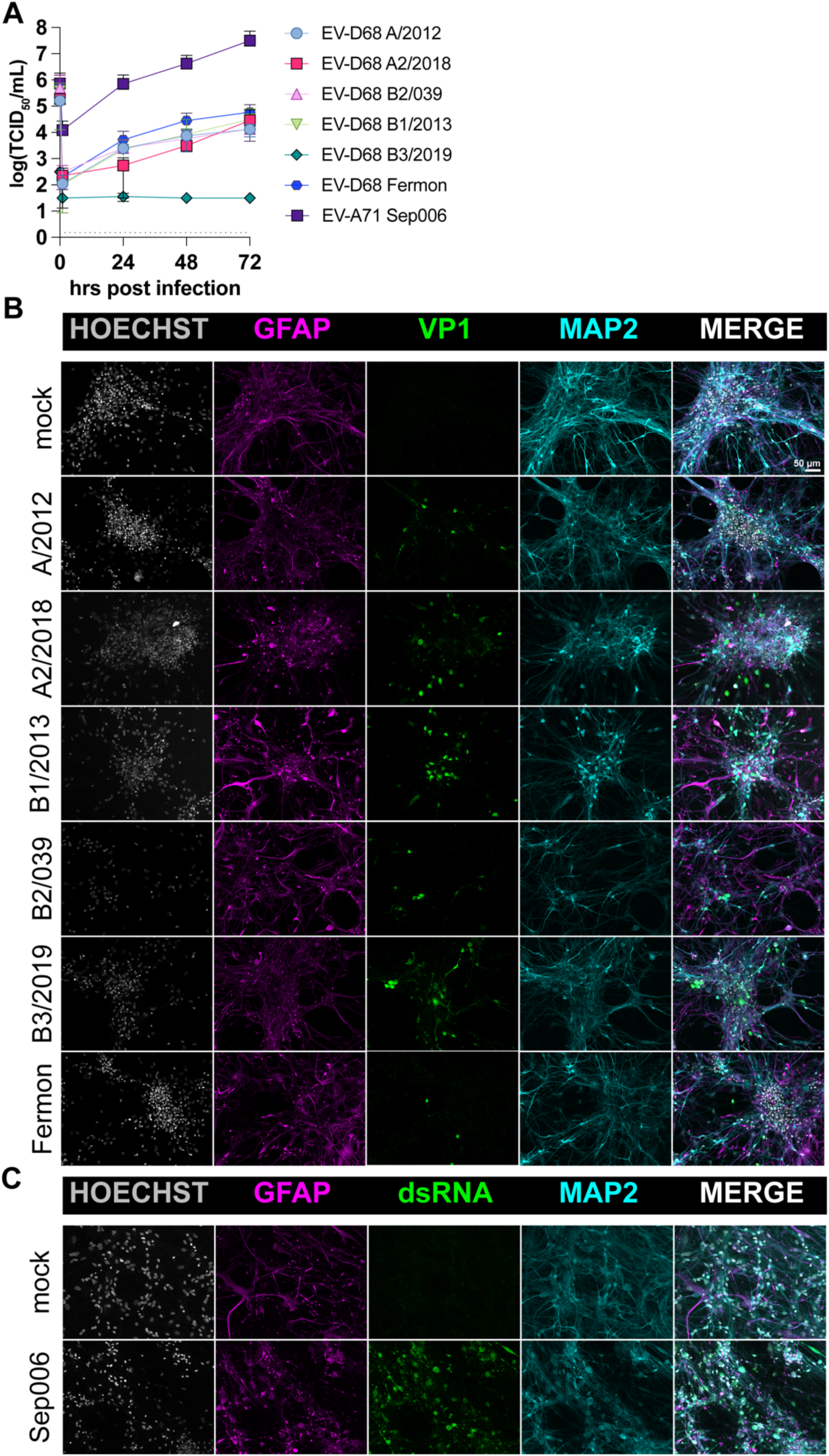
Enterovirus-D68 and Enterovirus-A71 productively replicate in co-cultures and show similarity in their cell tropism. (A) Neural co-cultures were infected with different EV-D68 strains and the EV-A71 strain Sep006 from the C4 genotype at a MOI of 0.1. Viral titres were determined in the supernatant at the indicated timepoints by endpoint dilution, with the first data point being the back-titration of the inoculum. Data represent mean ± SEM from three independent differentiation experiments and every growth curve was performed in triplicate. (B) At 24 hpi, the co-cultures were fixed and stained for the presence of EV-D68 structural antigen VP1 (green). MAP2 (cyan) was used as a marker for neurons, and astrocytes were identified by staining for GFAP (magenta). Cells were counterstained with Hoechst (grey) to visualize the nuclei. (C) At 24 hpi, EV-A71 inoculated co-cultures were fixed and stained for the presence of dsRNA, a marker for active EV-A71 replication (green), astrocytes (GFAP, magenta) and neurons (MAP2, cyan). The immunofluorescence data shown are representative examples from three independent experiments for each culture condition. Maximum intensity projections of Z-stacks are displayed. Abbreviations: EV = enterovirus; MOI = multiplicity of infection; SEM = standard error of the main; hpi = hours post inoculation; VP1 = viral protein 1; MAP2 = microtubule-associated protein 2; GFAP = glial fibrillary acidic protein; dsRNA = double-stranded RNA

Next, we wanted to investigate whether enterovirus infection induced apoptosis in GFAP^+^ astrocytes or MAP2^+^ neurons. Therefore, we inoculated with a higher MOI and stained for the apoptosis marker cleaved caspase-3 (CC-3) at 24 hpi or 72 hrs, respectively. We detected co-localization of CC-3^+^ and MAP2^+^ cells occasionally in mock or virus-inoculated cultures. Neither neurons nor astrocytes expressing dsRNA as a marker for virus replication showed co-localization with CC-3 similar to what we have observed with other viruses previously **(Fig. 3A and 3B).**^40^

**Figure 3.**
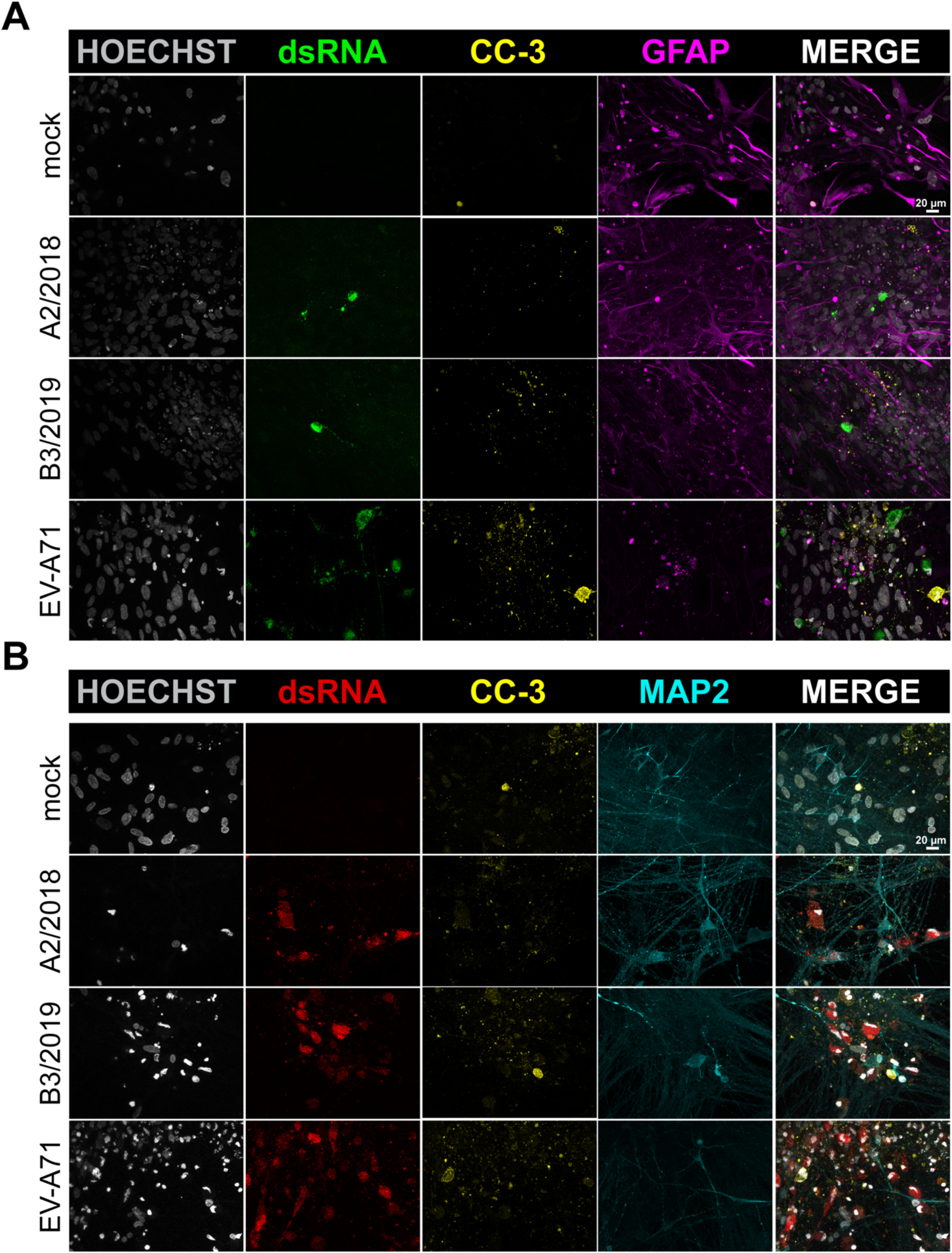
Enterovirus infection does not result in upregulation of cleaved caspase-3. Neural co-cultures were infected with EV-D68 or EV-A71 at an MOI of 1. At 24 and 72 hpi, the cells were fixed and stained for the presence dsRNA (green/red) as marker for virus infection, for either neural marker MAP2 (cyan) or astrocytic marker GFAP (magenta), and for the apoptosis marker CC-3 (yellow). Data shown are representative examples from two independent experiments. Abbreviations: EV = enterovirus; MOI = multiplicity of infection; hpi = hours post inoculation; VP1 = viral protein 1; dsRNA = double-stranded RNA; MAP2 = microtubule-associated protein 2; GFAP = glial fibrillary acidic protein; CC-3 = cleaved caspase-3

### Non-polio enterovirus infections exert pleiotropic negative effects on neural electrophysiology

To investigate the neurovirulent potential of currently circulating EV-D68 A2/2018 and B3/2019, as well as EV-A71 Sep006, we inoculated neural co-cultures and measured neural activity using a MEA system with an MOI of 1. We performed principal component analysis (PCA) to investigate patterns in our data obtained from the MEA recordings, enabling clear visualization and in-depth interpretation of complex, high-dimensional data (**Fig. 4; Supplementary Fig. 5**). To correct for potential virus-induced cell death, cell viability measurements were conducted before each recording. Additionally, we checked whether neural co-cultures had sufficient active electrodes and displayed similar spontaneous activity (**Fig. 5A; Supplementary Fig. 6; Supplementary Fig. 7**). At baseline, there were no significant differences observed between designated mock and inoculated group in the number of covered or active electrodes (**Fig. 5A; Supplementary Fig. 6A**). All groups displayed similar spontaneous activity represented by firing rate, burst frequency and network burst frequency at baseline before virus inoculation (**Supplementary Fig. 6A**).

**Figure 4.**
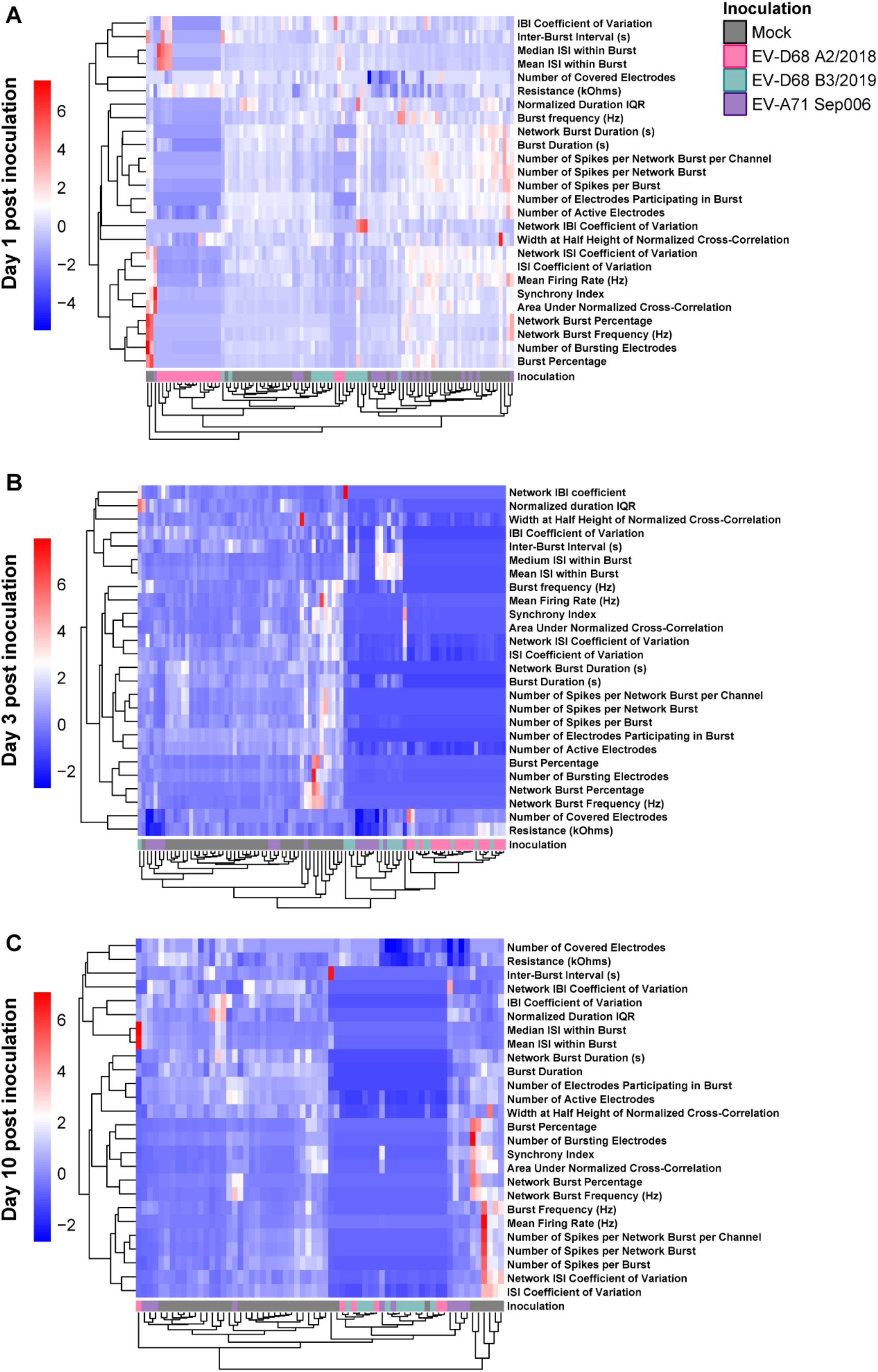
Heatmap of normalized neural activity data recorded from neural co-cultures inoculated with non-polio enteroviruses. Neural activity was recorded from neural co-cultures, consisting of Ngn2 neurons and astrocytes, on a MEA platform. Neural co-cultures were mock-inoculated or inoculated with EV-D68 A2/2018, B3/2019, or EV-A71 Sep006 with a MOI of 1. Only data that met inclusion criteria for MEA recordings were used (see Material and Methods). Any experiment that contained *Not Available* values throughout any of the chosen MEA output variables (Table 3) was excluded for PCA analysis, to ensure biological relevance and integrity of the dataset. Data was normalized against baseline recording obtained before inoculation. Data is shown from one (A), three (B), or 10 dpi (C). Rows represent output variables of MEA data and columns represent experimental data from a single well. All MEA data were z-score normalized for each variable across inoculation groups to enable cross-condition comparisons. Color intensity reflects a positive or negative effect on neural activity, with white to red indicating a more positive effect. Abbreviations: Ngn2 = Neurogenin-2; MEA = micro-electrode array; EV = enterovirus; MOI = multiplicity of infection; dpi = days post inoculation;

**Figure 5.**
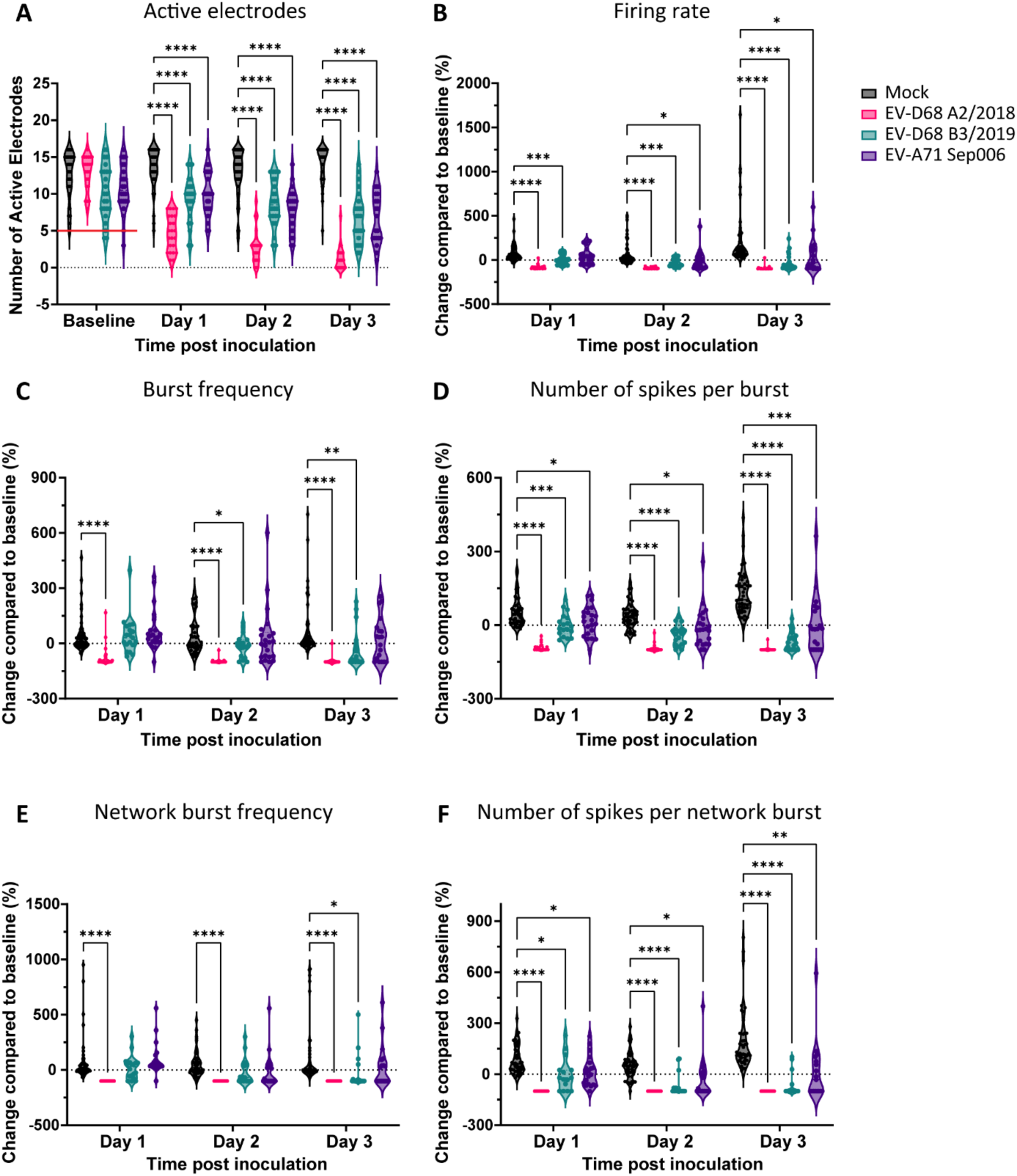
Enterovirus infection impacts the spontaneous activity of neural co-culture. Neural co-cultures were inoculated with EV-D68 A2/2018 (pink), B3/2019 (cyan), or EV-A71 Sep006 (purple) with a MOI of 1, and neural activity was measured between one to three dpi. The following parameters were displayed (A) number of active electrodes, where wells were excluded if they contained 5 or less active electrodes at baseline (indicated with a red line); (B) firing rate; (C) burst frequency, (D) number of spikes per burst, (E) network burst frequency and (F) number of spikes per network burst. Data displayed are derived from at least four independent experiments (*n* = 48 for control and *n* = 24 per inoculation group, see exclusion criteria in the Material and Methods section). Statistical significance was calculated with a two-way ANOVA with a Šídák’s multiple comparisons post hoc test. Asterisks indicate statistical significance (\**P*<0.05, \*\**P*<0.01, \*\*\**P*<0.001, \*\*\*\**P*<0.0001). Abbreviations: EV = enterovirus; MOI = multiplicity of infection; dpi = days post inoculation.

First, we investigated the hierarchical clustering of the output variables of the MEA and the key influencing parameters upon infection (**Fig. 4**). EV-D68 A2/2018 inoculated cultures clustered together and affected the output variables of the MEA with a mostly negative effect already 1 day post inoculation (dpi; **Fig. 4A**). The most negative effect was observed in variables such as “Burst Frequency,” “Burst Duration,” “Number of Active Electrodes,” “Number of Spikes per (Network) Burst,” indicating that neural activity is inhibited by EV-D68 A2/2018. PCA analysis confirmed that EV-D68 A2/2018 is more distinct from EV-D68 B3/2019, EV-A71 Sep006, or mock-inoculated controls, suggesting the strongest effect of EV-D68 A2/2018 in impairment of the neural network (**Supplementary Fig. 5A**). Over the course of infection, the two strains of EV-D68 occupied the same area in the PCA blots indicating a similar negative effect on the neural activity that is distant from EV-A71 at 3 dpi (**Fig. 4B and Supplementary Fig. 5B**) and 10 dpi (**Fig. 4C and Supplementary Fig. 5C**). In the mock cultures, a positive effect is observed on most of the MEA output variables, indicating that these cultures are in development and neural activity is increasing (**Fig. 4C and Fig. 1D**). Overall, PCA and heatmap analysis is suggestive that EV-D68 and EV-A71 infections result in adverse changes in the MEA outcomes to varying degrees.

### Non-polio enteroviruses infection reduces the network activity of hPSC-derived neural co-cultures

We measured spontaneous and network activity of neural co-cultures over ten days of infection with EV-D68 A2/2018 and B3/2019, and EV-A71 Sep006. A baseline recording of each individual well was performed before inoculation to compare the direct viral effect on the network. All three virus strains caused significant reductions in neural activity, but with variations in their temporal patterns. The number of covered electrodes decreased significantly for EV-A71 Sep006 starting at 2 days post-infection and continuing through day 10, while EV-D68 A2/2018 showed this effect from days 5-10, and EV-D68 B3/2019 only affected electrode coverage on days 9-10 (**Supplementary Fig. 6A**), possibly reflecting cell death upon infection. All three viruses caused significant decrease in active electrode number between days 1-3 post-infection (**Fig. 5A**), and these reductions persisted throughout the experimental period of 10 days (**Supplementary Fig. 6B**), This suggests that neural activity is inhibited upon virus-inoculation by a virus-induced mechanism. This is also shown when the covered electrodes are compared to the active electrodes (**Supplementary Fig. 6B**). EV-D68 A2/2018 inoculation has the strongest effect on the number of active electrodes (**Supplementary Fig. 6C**), followed by EV-D68 B3/2019 (**Supplementary Fig. 6D**). Both of the viruses cause cell death measured by the number of covered electrodes, yet with temporal differences. Meanwhile, EV-A71 Sep006 seems to follow a different pattern, where covered and active electrodes are more correlated to each other (**Supplementary Fig. 6E**). We further investigated different variables indicative for spontaneous neural activity. The neural firing rate was significantly reduced in cultures inoculated with EV-D68 A2/2018 and B3/2019 from days 1-3 and continued through day 10. In contrast, EV-A71 Sep006 inoculated cultures showed more variation, and decreased firing rates were observed between 2-3 dpi and 8-10 dpi. (**Fig. 5B, Supplementary Fig. 7C**). EV-D68 A2/2018 inoculated cultures showed significant decreases in burst frequency starting from day 1 onward, while EV-D68 B3/2019 inoculated cultures demonstrated this effect beginning from day 2 onward (**Fig. 5C; Supplementary Fig. 7D**). EV-A71 Sep006 did not alter the burst frequency (**Fig. 5C; Supplementary Fig. 7D**). All viruses significantly decreased the number of spikes per burst beginning at 1 dpi and persisting through 10 dpi (**Fig. 5D; Supplementary Fig. 7E**). Overall, these results indicate that while all three virus strains impair neural function, EV-D68 A2/2018 and B3/2019 strains produce more sustained and consistent neurotoxic effects across multiple parameters compared to EV-A71 Sep006.

To further characterize viral effects on neural network activity, we analyzed network burst parameters, including network burst frequency and spike density within bursts. Network burst frequency was significantly reduced in EV-D68 A2/2018 inoculated cultures beginning at 1 dpi, while in EV-D68 B3/2019 inoculated cultures this effect started 3 dpi and, in both cases, persisted until 10 dpi (**Fig. 5E; Supplementary Fig. 7F**). EV-A71 Sep006 did not alter the network burst frequency (**Fig. 5E; Supplementary Fig. S7F**). Analysis of spikes per network burst revealed that all three virus strains significantly decreased the number of spikes per burst during the early infection period (1-3 dpi) (**Fig. 5F**). However, the temporal patterns diverged during later time points. EV-D68 A2/2018 and B3/2019 inoculated cultures significantly reduced spike counts per burst throughout the entire 10-day observation period (**Supplementary Fig. 7G**). In contrast, EV-A71 Sep006 inoculated cultures showed significant reductions occurring only at 5 dpi and from 7-10 dpi. Taken together, these findings indicate that EV-D68 and EV-A71 inoculation disrupt coordinated network activity, and they exhibit distinct temporal profiles of network dysfunction. The EV-D68 strains A2/2018 and B3/2019 showed a more pronounced effect on inhibiting the neural network parameters compared to EV-A71. Even though the onset of inhibition differed between EV-D68 and EV-A71, the same parameters were affected over time.

## Discussion

The re-emerging non-polio enteroviruses EV-D68 and EV-A71 are major causes of viral encephalitis, yet their underlying neuropathogenesis remains poorly understood.^1,47,48^ In this study, we investigated the replication efficiency and cellular tropism of various EV-D68 and EV-A71 strains, as well as their impact on spontaneous activity, using a physiologically relevant human neural co-culture model consisting of hPSC-derived excitatory cortical Ngn2 neurons and astrocytes. Reduction of spontaneous activity and decreased network activity were more pronounced in EV-D68 inoculated cultures compared to EV-A71.

EV-D68 and EV-A71 showed slight differences in their cellular neurotropism. While EV-D68 preferentially infected MAP2^+^ neurons and only sporadically GFAP^+^ astrocytes, EV-A71 infected MAP2^+^ neurons and GFAP^+^ astrocytes equally efficiently. It has been shown that EV-D68 infects human astrocytes and mouse astrocytes,^49,50^ but not astrocytes in a primary rat neuronal model.^34^ EV-A71 has been detected in neurons, astrocytes, as well as neuron progenitor cells.^23–25,51^ EV-A71 and all EV-D68 viruses, except EV-D68 B3/2019, were able to replicate in neural co-cultures productively. This is in line with previous studies performed in primary rat cortical neurons, a neuron-like cell line and hPSC-derived forebrain and cerebral organoids.^34,50,52,53^ Similar to other studies, our data also confirm that the ability of different EV-D68 clades to replicate efficiently in neurons is not a recently acquired phenotype, as older isolates also replicate productively.^46,50,54^ Our results suggest that all EV-D68 strains are able to release viral progeny into the supernatants, except EV-D68 B3/2019 which was impaired in the release of infectious virus particles. However, in both cases, EV-D68 viruses were able to spread through the cultures over the course of the experiments. This potentially indicates differences in the spreading or release mechanism between EV-D68 strains (e.g. transsynaptic spread or extracellular vesicles release).^55^ Whether this affects the neurovirulent potential of these viruses is currently not understood.

The reduction in the number of covered electrodes suggests that all three viruses induced cell death and EV-D68 A2/2018 showed the strongest phenotype compared to EV-D68 B3/2019 and EV-A71. Which type of cell death is induced by these pathogens in our cultures and whether they differ between the viruses remains to be established. It has been shown that EV-A71 is able to induce cell death through apoptosis, pyroptosis, and ferroptosis in different neural cultures,^56–60^ while EV-D68 induced a non-apoptotic necrosis-like cell death.^34^ In our model, we did not observe induction of cleaved caspase-3 upon inoculation with EV-D68 or EV-A71. Neuronal cell death can result from the impairment of neurotransmission or be directly induced by infection. It is interesting to note, that inoculation with EV-A71 did not enhance cell death despite a broader cell tropism of this virus, that could potentially result in virus-induced dysfunction of both cell types in our cultures. It remains plausible that the observed cell death is a consequence of neural activity inhibition or of an alternative active or passive cell death pathway, distinct from apoptosis, triggered upon inoculation.

EV infection affected both spontaneous neuronal activity and overall neural network function. EV-D68 A2/2019 induced a rapid shut-down of neural activity. EV-D68 B3/2019 followed this trend, although more slowly and less extensively. Despite the significant differences in the replication between EV-A2/2018 and B3/2019, both viruses produced a sustained and consistent neurotoxic effect. This aligns with earlier research, which found that different EV-D68 viruses are causing a persistent reduction in neural activity in rat primary neurons.^34^ In comparison, EV-A71 Sep006 showed a slower temporal kinetics in impairing neural activity, despite more efficient replication in the neural co-cultures. This suggests that the neurotoxic effects are not determined by viral load but are rather virus-specific. Further studies comparing changes in gene expression or metabolomics can help shed light on the mechanisms responsible for the decrease in neurotransmission for EV-D68 and EV-A71.^61,62^

Our hPSC-derived neural co-culture model provides a valuable and scalable platform for studying neuronal activity during viral infection and other brain disorders.^63,64^ However, it does not fully recapitulate the cellular diversity, structural organization, immune interactions, and network complexity of the human brain. For example, our model does not contain any inhibitory neurons. Yet, we do observe similar outcomes as in a primary rat cortical neurons model where both excitatory and inhibitory neurons were infected with EV-D68.^34^ Additionally, because of the lack of immune competence in our model, we cannot investigate the full range of brain inflammatory response as reported in EV-induced human encephalitis cases and rat neuronal cells.^24,26,27,60^ Which inflammatory responses contribute to the development of EV-induced encephalitis and whether this differs between enterovirus species is currently unknown.

Our study demonstrates that EV infections have profound effects on the neural network activity providing a link between EV infection and disturbances in the brain homeostasis and functionality using a physiologically relevant hPSC-derived neural *in vitro* model. Neurotropic mechanisms leading to this effect are virus-specific and can include dysregulation of neural homeostasis, cell death, inflammatory responses and changes in neuroplasticity. Integration of electrophysiological outcomes presented here with metabolomics and gene expression/omics data in future studies holds great promise in understanding the neuropathogenesis of enteroviruses on the cellular level.

## Acknowledgements

We thank Adam Meijer from the National Institute for Public Health and Environment (RIVM) in the Netherlands for providing the EV-D68 isolates. We thank Dr. Stefan Barakat (Department of Clinical Genetics Erasmus MC University Medical Center, Rotterdam, the Netherlands) for carefully reading the manuscript and providing suggestions for improvements. We thank David den Ouden for scientific support.

## Authors Contribution

Conceptualization FB, DVR, KL, LB

Investigation FB, SS, KL, LB

Formal Analysis FB, SS, MP, KL, LB

Resources BM, WR, FMSDV, DvR, LB

Methodology FB, WR, FMSDV, KL, LB

Supervision LB

Visualization FB, MP, KL, LB

Writing Original FB, KL, LB

Writing-Reviewing all authors

Funding acquisition FMSDV, DVR KL, LB

## References

1. Harvala H, Johannesen CK, Benschop KSM, et al. Enterovirus circulation in the WHO European region, 2015-2022: a comparison of data from WHO’s three core poliovirus surveillance systems and the European Non-Polio Enterovirus Network (ENPEN). Lancet Reg Health Eur. 2025;53:101292. doi:10.1016/j.lanepe.2025.101292

2. Rankin DA, Spieker AJ, Perez A, et al. Circulation of Rhinoviruses and/or Enteroviruses in Pediatric Patients With Acute Respiratory Illness Before and During the COVID-19 Pandemic in the US. JAMA Netw Open. 2023;6(2):e2254909. doi:10.1001/jamanetworkopen.2022.54909

3. Nguyen-Tran H, Butler M, Simmons D, Dominguez SR, Messacar K. Return of the Biennial Circulation of Enterovirus D68 in Colorado Children in 2024 Following the Large 2022 Outbreak. Viruses. 2025;17(5):673. doi:10.3390/v17050673

4. de Schrijver S, Vanhulle E, Ingenbleek A, et al. Epidemiological and clinical insights into enterovirus circulation in Europe, 2018 - 2023: a multi-center retrospective surveillance study. J Infect Dis. Published online April 4, 2025:jiaf179. doi:10.1093/infdis/jiaf179

5. Chuang YY, Huang YC. Enteroviral infection in neonates. J Microbiol Immunol Infect. 2019;52(6):851–857. doi:10.1016/j.jmii.2019.08.018

6. Hawkes MT, Vaudry W. Nonpolio enterovirus infection in the neonate and young infant. Paediatr Child Health. 2005;10(7):383–388.

7. Cassidy H, van Leer-Buter C, Niesters HGM. Enterovirus Infections in Solid Organ Transplant Recipients: a Clinical Comparison from a Regional University Hospital in the Netherlands. Microbiol Spectr. 10(1):e02215–21. doi:10.1128/spectrum.02215-21

8. Grizer CS, Messacar K, Mattapallil JJ. Enterovirus-D68 - A Reemerging Non-Polio Enterovirus that Causes Severe Respiratory and Neurological Disease in Children. Front Virol. 2024;4:1328457. doi:10.3389/fviro.2024.1328457

9. Jartti M, Flodström-Tullberg M, Hankaniemi MM. Enteroviruses: epidemic potential, challenges and opportunities with vaccines. J Biomed Sci. 2024;31(1):73. doi:10.1186/s12929-024-01058-x

10. Xie Z, Khamrin P, Maneekarn N, Kumthip K. Epidemiology of Enterovirus Genotypes in Association with Human Diseases. Viruses. 2024;16(7):1165. doi:10.3390/v16071165

11. Whitehouse ER, Lopez A, English R, et al. Surveillance for Acute Flaccid Myelitis - United States, 2018-2022. MMWR Morb Mortal Wkly Rep. 2024;73(4):70-76. doi:10.15585/mmwr.mm7304a1

12. Messacar K, Spence-Davizon E, Osborne C, et al. Clinical characteristics of enterovirus A71 neurological disease during an outbreak in children in Colorado, USA, in 2018: an observational cohort study. Lancet Infect Dis. 2020;20(2):230-239. doi:10.1016/S1473-3099(19)30632-2

13. Bosch A, Carcereny A, García-Pedemonte D, et al. Human enteroviruses and the long road to acute flacid paralysis eradication. J Appl Microbiol. 2025;136(1):lxae311. doi:10.1093/jambio/lxae311

14. Puenpa J, Wanlapakorn N, Vongpunsawad S, Poovorawan Y. The History of Enterovirus A71 Outbreaks and Molecular Epidemiology in the Asia-Pacific Region. J Biomed Sci. 2019;26(1):75. doi:10.1186/s12929-019-0573-2

15. Messali S, Bertelli A, Dotta L, et al. Outbreak of Enterovirus D68 in Young Children, Brescia, Italy, August to November 2024. J Med Virol. 2025;97(5):e70372. doi:10.1002/jmv.70372

16. Fall A, Norton JM, Abdullah O, Pekosz A, Klein E, Mostafa HH. Enhanced genomic surveillance of enteroviruses reveals a surge in enterovirus D68 cases, the Johns Hopkins health system, Maryland, 2024. J Clin Microbiol. 63(7):e00469-25. doi:10.1128/jcm.00469-25

17. Hirvonen A, Johannesen CK, Simmonds P, et al. Sustained circulation of enterovirus D68 in Europe in 2023 and the continued evolution of enterovirus D68 B3-lineages associated with distinct amino acid substitutions in VP1 protein. J Clin Virol. 2025;178:105785. doi:10.1016/j.jcv.2025.105785

18. Freeman MC, Messacar K. Enterovirus and Parechovirus Neurologic Infections in Children: Clinical Presentations and Neuropathogenesis. J Pediatric Infect Dis Soc. 2025;14(1):piae069. doi:10.1093/jpids/piae069

19. Volle R, Archimbaud C, Couraud PO, et al. Differential permissivity of human cerebrovascular endothelial cells to enterovirus infection and specificities of serotype EV-A71 in crossing an in vitro model of the human blood–brain barrier. Journal of General Virology. 2015;96(7):1682–1695. doi:10.1099/vir.0.000103

20. Gaume L, Chabrolles H, Bisseux M, et al. Enterovirus A71 crosses a human blood-brain barrier model through infected immune cells. Microbiol Spectr. 2024;12(6):e0069024. doi:10.1128/spectrum.00690-24

21. Laksono BM, Sooksawasdi Na Ayudhya S, Aguilar-Bretones M, Embregts CWE, van Nierop GP, van Riel D. Human B cells and dendritic cells are susceptible and permissive to enterovirus D68 infection. mSphere. 2024;9(2):e0052623. doi:10.1128/msphere.00526-23

22. Maloney JA, Mirsky DM, Messacar K, Dominguez SR, Schreiner T, Stence NV. MRI findings in children with acute flaccid paralysis and cranial nerve dysfunction occurring during the 2014 enterovirus D68 outbreak. AJNR Am J Neuroradiol. 2015;36(2):245–250. doi:10.3174/ajnr.A4188

23. Li J, Chen F, Liu T, Wang L. MRI findings of neurological complications in hand-foot-mouth disease by enterovirus 71 infection. Int J Neurosci. 2012;122(7):338–344. doi:10.3109/00207454.2012.657379

24. Wong KT, Munisamy B, Ong KC, et al. The distribution of inflammation and virus in human enterovirus 71 encephalomyelitis suggests possible viral spread by neural pathways. J Neuropathol Exp Neurol. 2008;67(2):162–169. doi:10.1097/nen.0b013e318163a990

25. Feng M, Guo S, Fan S, et al. The Preferential Infection of Astrocytes by Enterovirus 71 Plays a Key Role in the Viral Neurogenic Pathogenesis. Front Cell Infect Microbiol. 2016;6:192. doi:10.3389/fcimb.2016.00192

26. Vogt MR, Wright PF, Hickey WF, De Buysscher T, Boyd KL, Crowe JE. Enterovirus D68 in the Anterior Horn Cells of a Child with Acute Flaccid Myelitis. N Engl J Med. 2022;386(21):2059–2060. doi:10.1056/NEJMc2118155

27. Yang Y, Wang H, Gong E, et al. Neuropathology in 2 cases of fatal enterovirus type 71 infection from a recent epidemic in the People’s Republic of China: a histopathologic, immunohistochemical, and reverse transcription polymerase chain reaction study. Hum Pathol. 2009;40(9):1288–1295. doi:10.1016/j.humpath.2009.01.015

28. Chang LY, Huang LM, Gau SSF, et al. Neurodevelopment and cognition in children after enterovirus 71 infection. N Engl J Med. 2007;356(12):1226–1234. doi:10.1056/NEJMoa065954

29. Li Y, Yang J, Liang L, et al. Clinical characteristics and severity of hand, foot, and mouth disease by virus serotype: A prospective hospital-based cohort study. PLoS Negl Trop Dis. 2025;19(5):e0013039. doi:10.1371/journal.pntd.0013039

30. Hosseini S, Wilk E, Michaelsen-Preusse K, et al. Long-Term Neuroinflammation Induced by Influenza A Virus Infection and the Impact on Hippocampal Neuron Morphology and Function. J Neurosci. 2018;38(12):3060–3080. doi:10.1523/JNEUROSCI.1740-17.2018

31. Gu L, Zhou Y, Wang G, et al. Spatial learning and memory impaired after infection of non-neurotropic influenza virus in BALB/c male mice. Biochemical and Biophysical Research Communications. 2021;540:29–36. doi:10.1016/j.bbrc.2020.12.092

32. Jiang Q, Li G, Wang H, et al. SARS-CoV-2 spike S1 protein induces microglial NLRP3-dependent neuroinflammation and cognitive impairment in mice. Exp Neurol. 2025;383:115020. doi:10.1016/j.expneurol.2024.115020

33. Park H, Yu JE, Kim S, Nahm SS, Chung C. Decreased Na(+) influx lowers hippocampal neuronal excitability in a mouse model of neonatal influenza infection. Sci Rep. 2015;5:13440. doi:10.1038/srep13440

34. Poelaert KCK, van Kleef RGDM, Liu M, et al. Enterovirus D-68 Infection of Primary Rat Cortical Neurons: Entry, Replication, and Functional Consequences. mBio. 2023;14(2):e0024523. doi:10.1128/mbio.00245-23

35. Frega M, van Gestel SHC, Linda K, et al. Rapid Neuronal Differentiation of Induced Pluripotent Stem Cells for Measuring Network Activity on Micro-electrode Arrays. J Vis Exp. 2017;(119). doi:10.3791/54900

36. Zhang Y, Pak C, Han Y, et al. Rapid Single-Step Induction of Functional Neurons from Human Pluripotent Stem Cells. Neuron. 2013;78(5):785–798. doi:10.1016/j.neuron.2013.05.029

37. Lendemeijer B, Unkel M, Smeenk H, et al. Human Pluripotent Stem Cell-Derived Astrocyte Functionality Compares Favorably with Primary Rat Astrocytes. eNeuro. 2024;11(9):ENEURO.0148-24.2024. doi:10.1523/ENEURO.0148-24.2024

38. Gunhanlar N, Shpak G, van der Kroeg M, et al. A simplified protocol for differentiation of electrophysiologically mature neuronal networks from human induced pluripotent stem cells. Mol Psychiatry. 2018;23(5):1336–1344. doi:10.1038/mp.2017.56

39. Shih PY, Kreir M, Kumar D, et al. Development of a fully human assay combining NGN2-inducible neurons co-cultured with iPSC-derived astrocytes amenable for electrophysiological studies. Stem Cell Research. 2021;54:102386. doi:10.1016/j.scr.2021.102386

40. Bauer L, Lendemeijer B, Leijten L, et al. Replication Kinetics, Cell Tropism, and Associated Immune Responses in SARS-CoV-2- and H5N1 Virus-Infected Human Induced Pluripotent Stem Cell-Derived Neural Models. mSphere. 2021;6(3):e0027021. doi:10.1128/mSphere.00270-21

41. REED LJ, MUENCH H. A SIMPLE METHOD OF ESTIMATING FIFTY PER CENT ENDPOINTS12. Am J Epidemiol. 1938;27(3):493–497. doi:10.1093/oxfordjournals.aje.a118408

42. Pré D, Wooten AT, Biesmans S, et al. Development of a platform to investigate long-term potentiation in human iPSC-derived neuronal networks. Stem Cell Reports. 2022;17(9):2141–2155. doi:10.1016/j.stemcr.2022.07.012

43. Bauer L, Manganaro R, Zonsics B, et al. Rational design of highly potent broad-spectrum enterovirus inhibitors targeting the nonstructural protein 2C. PLoS Biol. 2020;18(11):e3000904. doi:10.1371/journal.pbio.3000904

44. Paiva A, Park I, Principe J. A comparison of binless spike train measures. Neural Computing and Applications. 2010;19:405–419. doi:10.1007/s00521-009-0307-6

45. Pereirinha da Silva AK, van Trijp JP, Montenarie A, et al. Sialic Acid-Containing Glycolipids Extend the Receptor Repertoire of Enterovirus-D68. ACS Infect Dis. Published online July 9, 2025. doi:10.1021/acsinfecdis.5c00063

46. Hixon AM, Clarke P, Tyler KL. Contemporary Circulating Enterovirus D68 Strains Infect and Undergo Retrograde Axonal Transport in Spinal Motor Neurons Independent of Sialic Acid. J Virol. 2019;93(16):e00578–19. doi:10.1128/JVI.00578-19

47. Wang YC, Wu PH, Ting WC, et al. Single-cell signaling network profiling during redox stress reveals dynamic redox regulation in immune cells. Nat Commun. 2025;16(1):5600. doi:10.1038/s41467-025-60727-z

48. Halawani A, Khan S, Masood S, Firoze S. Enterovirus-Associated Meningoencephalitis and Enteroviruses in Patients with Acute Encephalitis. In: Sami H, Firoze S, Khan PA, eds. Viral and Fungal Infections of the Central Nervous System: A Microbiological Perspective. Springer Nature; 2023:97–123. doi:10.1007/978-981-99-6445-1_6

49. Zhang L, Yang J, Li H, et al. Enterovirus D68 Infection Induces TDP-43 Cleavage, Aggregation, and Neurotoxicity. J Virol. 2023;97(4):e0042523. doi:10.1128/jvi.00425-23

50. Rosenfeld AB, Warren AL, Racaniello VR. Neurotropism of Enterovirus D68 Isolates Is Independent of Sialic Acid and Is Not a Recently Acquired Phenotype. mBio. 2019;10(5):e02370–19. doi:10.1128/mBio.02370-19

51. Huang HI, Lin JY, Chen HH, et al. Enterovirus 71 infects brain-derived neural progenitor cells. Virology. 2014;468–470:592-600. doi:10.1016/j.virol.2014.09.017

52. Vazquez C, Negatu SG, Bannerman CD, Sriram S, Ming GL, Jurado KA. Antiviral immunity within neural stem cells distinguishes Enterovirus-D68 strain differences in forebrain organoids. J Neuroinflammation. 2024;21(1):288. doi:10.1186/s12974-024-03275-5

53. Sridhar A, Depla JA, Mulder LA, et al. Enterovirus D68 Infection in Human Primary Airway and Brain Organoids: No Additional Role for Heparan Sulfate Binding for Neurotropism. Microbiol Spectr. 2022;10(5):e0169422. doi:10.1128/spectrum.01694-22

54. Brown DM, Hixon AM, Oldfield LM, et al. Contemporary Circulating Enterovirus D68 Strains Have Acquired the Capacity for Viral Entry and Replication in Human Neuronal Cells. mBio. 2018;9(5):e01954–18. doi:10.1128/mBio.01954-18

55. Mutsafi Y, Altan-Bonnet N. Enterovirus Transmission by Secretory Autophagy. Viruses. 2018;10(3):139. doi:10.3390/v10030139

56. You L, Chen J, Liu W, et al. Enterovirus 71 induces neural cell apoptosis and autophagy through promoting ACOX1 downregulation and ROS generation. Virulence. 2020;11(1):537–553. doi:10.1080/21505594.2020.1766790

57. Chen TC, Lai YK, Yu CK, Juang JL. Enterovirus 71 triggering of neuronal apoptosis through activation of Abl-Cdk5 signalling. Cell Microbiol. 2007;9(11):2676–2688. doi:10.1111/j.1462-5822.2007.00988.x

58. Zhu X, Wu T, Chi Y, et al. Pyroptosis induced by enterovirus A71 infection in cultured human neuroblastoma cells. Virology. 2018;521:69–76. doi:10.1016/j.virol.2018.05.025

59. Chooi WH, Winanto null, Zeng Y, et al. Enterovirus-A71 preferentially infects and replicates in human motor neurons, inducing neurodegeneration by ferroptosis. Emerg Microbes Infect. 2024;13(1):2382235. doi:10.1080/22221751.2024.2382235

60. Chang CY, Li JR, Ou YC, et al. Enterovirus 71 infection caused neuronal cell death and cytokine expression in cultured rat neural cells. IUBMB Life. 2015;67(10):789–800. doi:10.1002/iub.1434

61. Capendale PE, García-Rodríguez I, Ambikan AT, et al. Parechovirus infection in human brain organoids: host innate inflammatory response and not neuro-infectivity correlates to neurologic disease. Nat Commun. 2024;15(1):2532. doi:10.1038/s41467-024-46634-9

62. Capendale PE, Ambikan AT, García-Rodríguez I, et al. Parechovirus-3 infection disrupts immunometabolism and leads to glutamate excitotoxicity in neural organoids. Published online September 14, 2024:2024.09.10.611955. doi:10.1101/2024.09.10.611955

63. Pradeepan KS, McCready FP, Wei W, et al. Calcium-Dependent Hyperexcitability in Human Stem Cell-Derived Rett Syndrome Neuronal Networks. Biol Psychiatry Glob Open Sci. 2024;4(2):100290. doi:10.1016/j.bpsgos.2024.100290

64. Frega M, Linda K, Keller JM, et al. Neuronal network dysfunction in a model for Kleefstra syndrome mediated by enhanced NMDAR signaling. Nat Commun. 2019;10(1):4928. doi:10.1038/s41467-019-12947-3

