## Supplementary Figures for "Non-polio enteroviruses compromise the electrophysiology of a human iPSC-derived neural network"

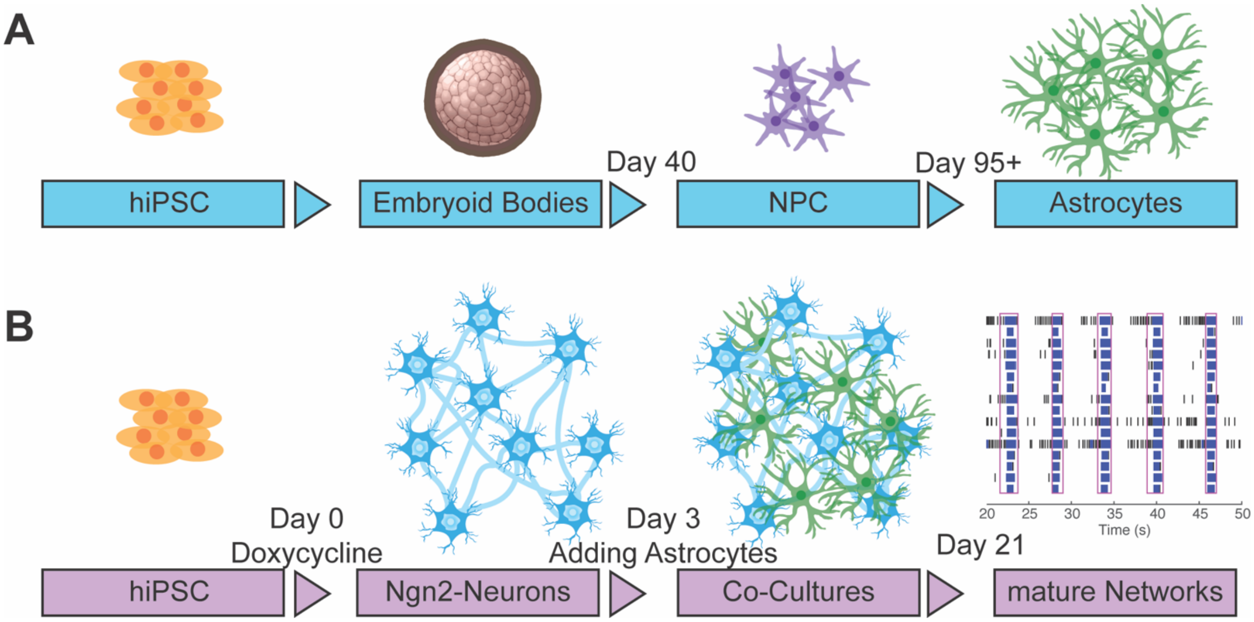


**Supplementary Figure 1. Overview of differentiation strategy.** (A) hiPSC are differentiated into NPCs and further into a pure culture of astrocytes as previously described1. (B) hiPSCs are differentiated into excitatory neurons by inducing overexpression of Ngn2 with Doxycycline. At day 3, astrocytes are added to the Ngn2-neurons in a 1:1 ratio and matured until DIV 21. From DIV 21, the Ngn2 neural co-cultures are electrophysiological active and used in experiments. Abbreviations: hiPSC = human induced pluripotent stem cells; NPC = neuron progenitor cells; Ngn2 = Neurogenin2; DIV = days in vitro;

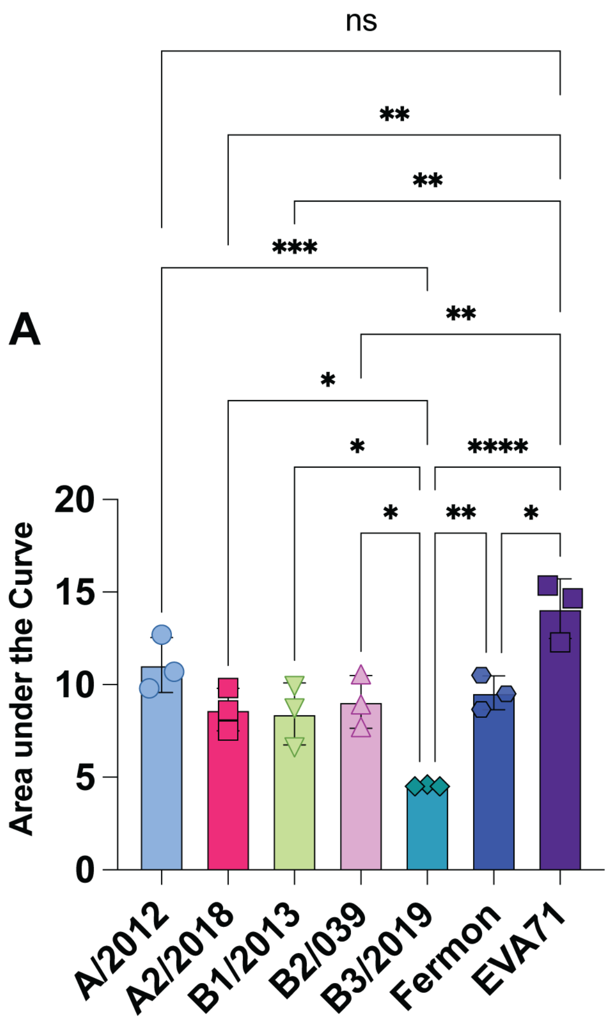


**Supplementary Figure 2. Area under the curve analysis of titres produced by Enterovirus-D68 and Enterovirus-A71 viruses in neural co-cultures.** (A) To evaluate statistical significances between the replication kinetics of EV-D68 and EV-A71 viruses in neural co-cultures, the area under the curve was calculated. The area under the curve for EV-A71 was correct for the higher input of EV-A71. Data displayed represent mean with standard deviation from three independent experiments. Differences in the growth curve between the different viruses were analyzed statistically with a One-Way Anova with a Tukey's post-hoc test to compare different groups.

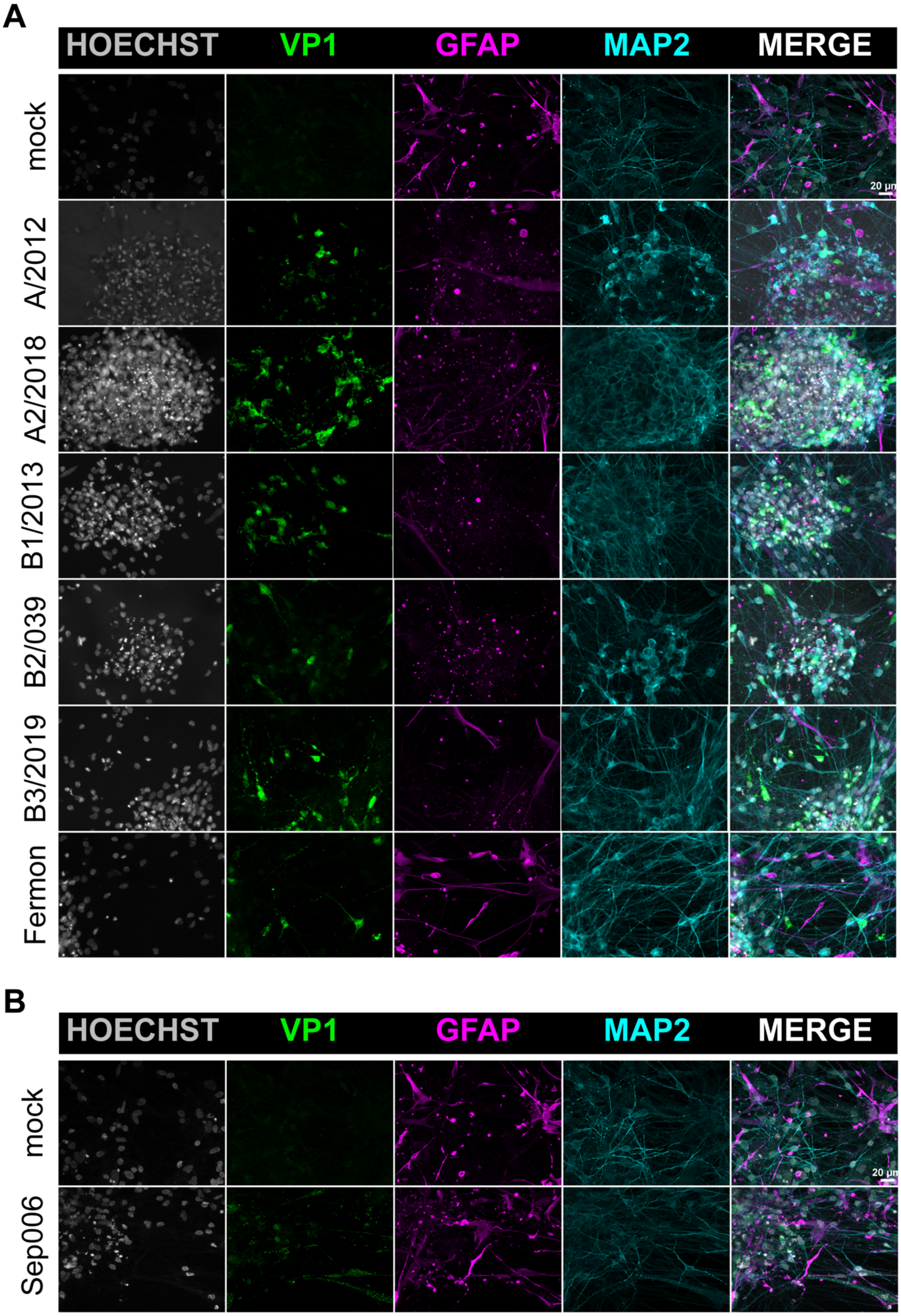
A
HOECHST
VP1
GFAP
MAP2
MERGE
mock
A/2012
A2/2018
B1/2013
B2/039
B3/2019
Fermon
20 µm
B
HOECHST
VP1
GFAP
MAP2
MERGE
mock
Sep006
20 µm

**Supplementary Figure 3. Enterovirus-D68 and Enterovirus-A71 infect neural co-cultures and show similarity in their cell tropism.** (A) 72 hpi, the neural co-cultures were fixed and stained for the presence of EV-D68 structural antigen VP1 (green). MAP2 (cyan) was used as a marker for neurons, astrocytes were identified by staining for GFAP (magenta). Cells were counterstained with Hoechst (grey) to visualize the nuclei. (B) 72 hpi, the co-cultures were fixed and stained for the presence of double stranded RNA, a marker for active EV-A71 replication (green), the astrocytic marker GFAP, neuron marker MAP2 and Hoechst. Immunofluorescence data shown are representative examples from three independent experiments. Maximum intensity projections of Z-stacks are displayed. Abbreviations: hpi = hours post inoculation; EV = enterovirus; VP1 = viral protein 1; MAP2 = microtubule-associated protein 2; GFAP = glial fibrillary acidic protein; dsRNA = double-stranded RNA

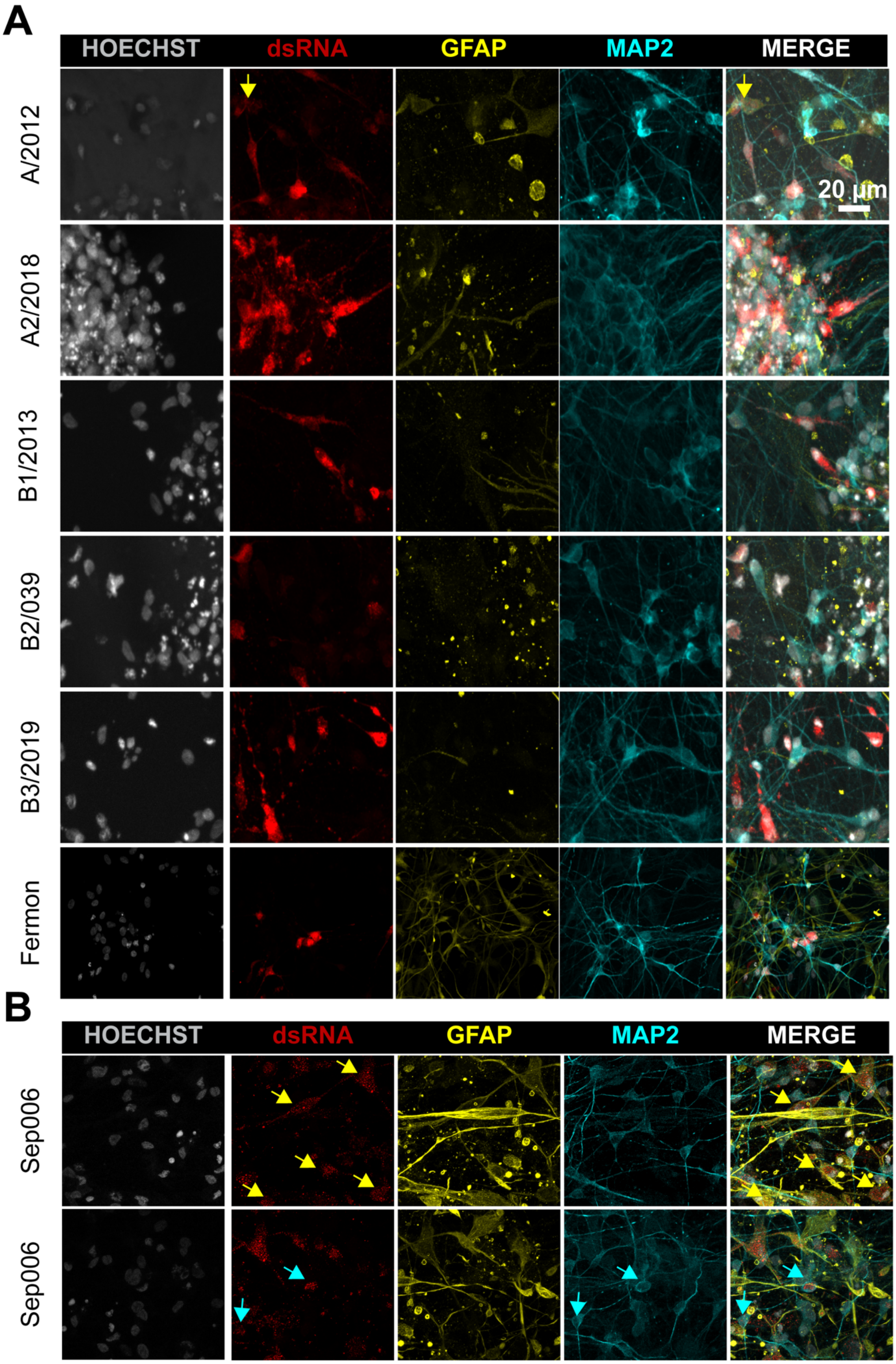
A
HOECHST
dsRNA
GFAP
MAP2
MERGE
A/2012
A2/2018
B1/2013
B2/039
B3/2019
Fermon
20 µm
B
HOECHST
dsRNA
GFAP
MAP2
MERGE
Sep006
Sep006

**Supplementary Figure 4. Cell tropism of EV-D68 and EV-A71 in neural co-cultures.** (A) 72 hpi, the neural co-cultures were fixed and stained for the presence of EV-D68 structural antigen VP1 (red). MAP2 (cyan) was used as a marker for neurons, astrocytes were identified by staining for GFAP (yellow). Cells were counterstained with Hoechst (grey) to visualize the nuclei. Yellow arrows represent infected astrocytes (B) 72 hpi, the co-cultures were fixed and stained for the presence of double stranded RNA, a marker for active EV-A71 replication (red), the astrocytic marker GFAP (yellow), neuron marker MAP2 (cyan) and Hoechst (grey). Yellow or cyan arrows represent infected astrocytes or neurons respectively.

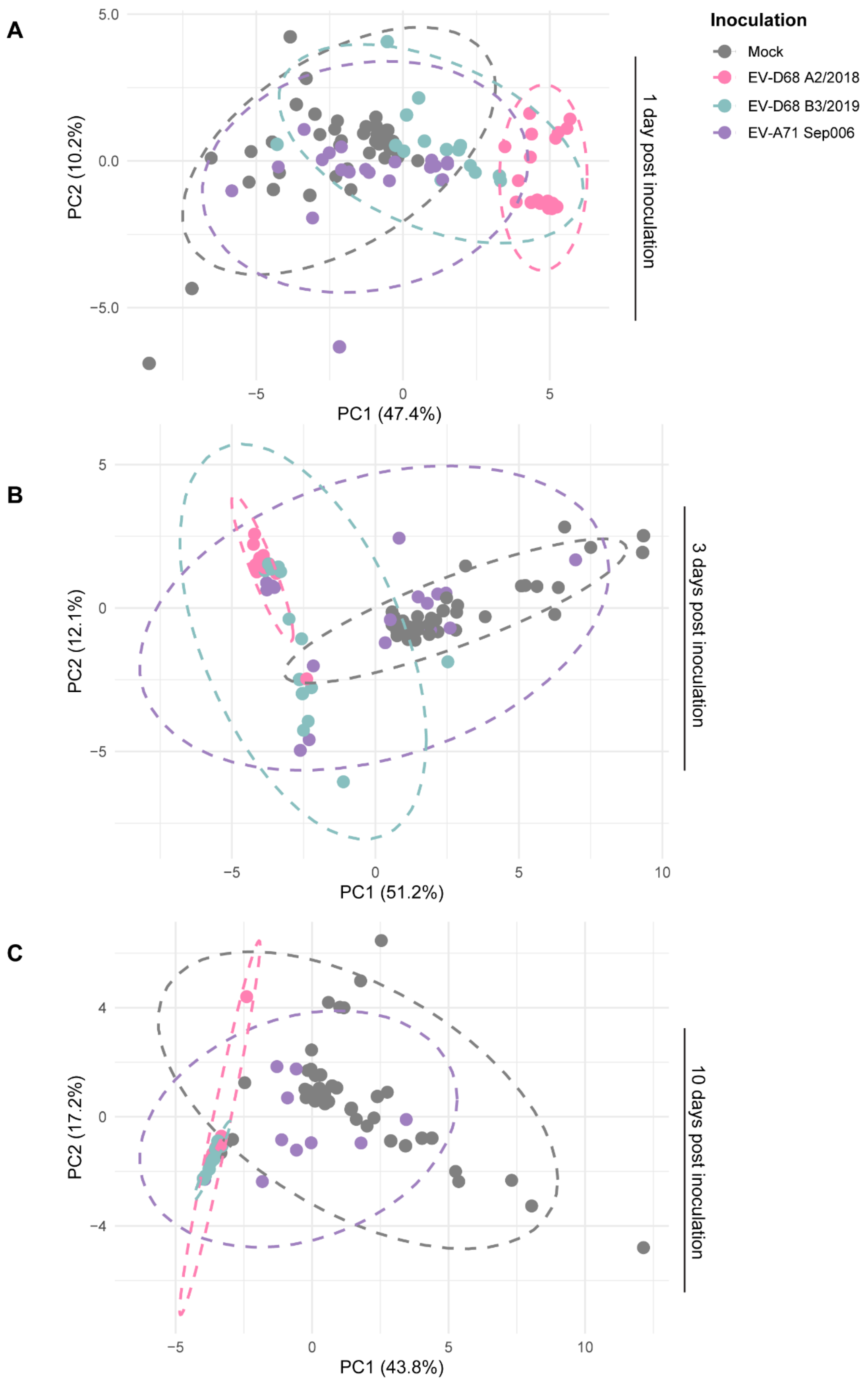

A
B
C
Inoculation
Mock
EV-D68 A2/2018
EV-D68 B3/2019
EV-A71 Sep006
PC1 (47.4%)
PC2 (10.2%)
1 day post inoculation
PC1 (51.2%)
PC2 (12.1%)
3 days post inoculation
PC1 (43.8%)
PC2 (17.2%)
10 days post inoculation

**Supplementary Figure 5. Principal Component Analysis (PCA) of neural activity recorded from co-cultures inoculated with Enterovirus-D68 A2, B3 or Enterovirus-A71 Sep006.** PCA was performed on neural data recorded from neural co-cultures, consisting of Ngn2 neurons and astrocytes on a MEA platform. Cultures were mock-inoculated, or with EV-D68 A2/2018, B3/2019 or EV-A71 Sep006 ($n$ = 48 for control and $n$ = 24 per inoculation group, unless datapoints were excluded based on exclusion criteria as described in the Material and Methods section). PCA plots of one (A), three (B), and ten dpi (C) are shown. The first two principal components (PC1 and PC2), accounting for 47.4% and 10.2% (1 dpi), 51.2% and 12.1% (3 dpi), and 43.8% and 17.2% (10 dpi) of the total variance, respectively, are shown. Each point represents experimental data from a single well, with colors indicating inoculation groups. PCA was conducted on all output variables of MEA data stated in the Material and Methods section. A 95% prediction ellipse is depicted for each group, highlighting differences in neural activity patterns across inoculation groups. Abbreviations: PCA = principal component analysis; Ngn2 = Neurogenin-2; MEA = micro-electrode array; EV = enterovirus; dpi = days post inoculation;

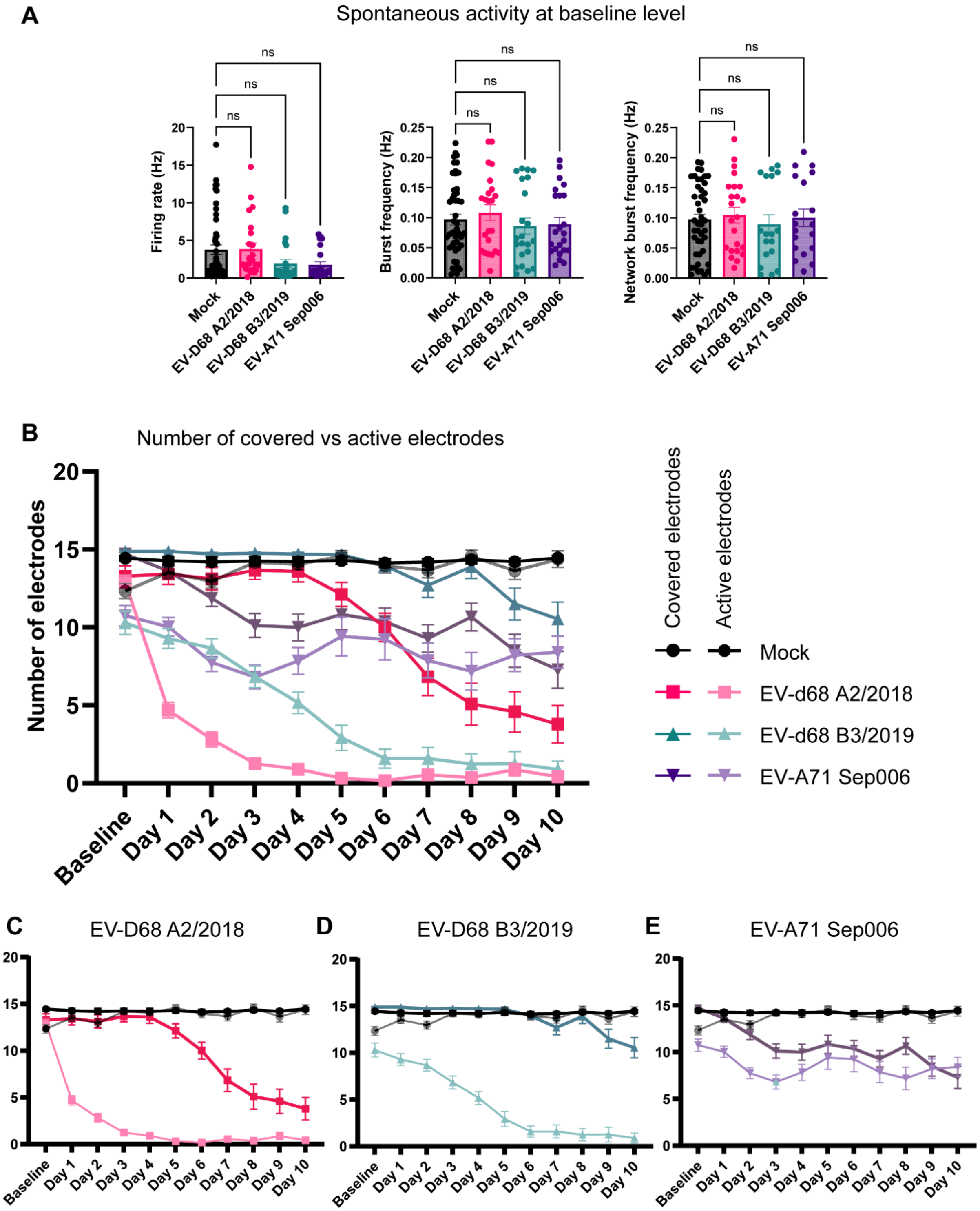


**Supplementary Figure 6. Neural Network activity of cultures. (A)** Firing rate, burst frequency, and network burst frequency at baseline recording. (B) Comparison of covered and active electrodes during the time of infection with the viruses (C) EV-D68 A2/2018, (D) EV-D68 B3/2019 and (E) EV-A71 Sep006.

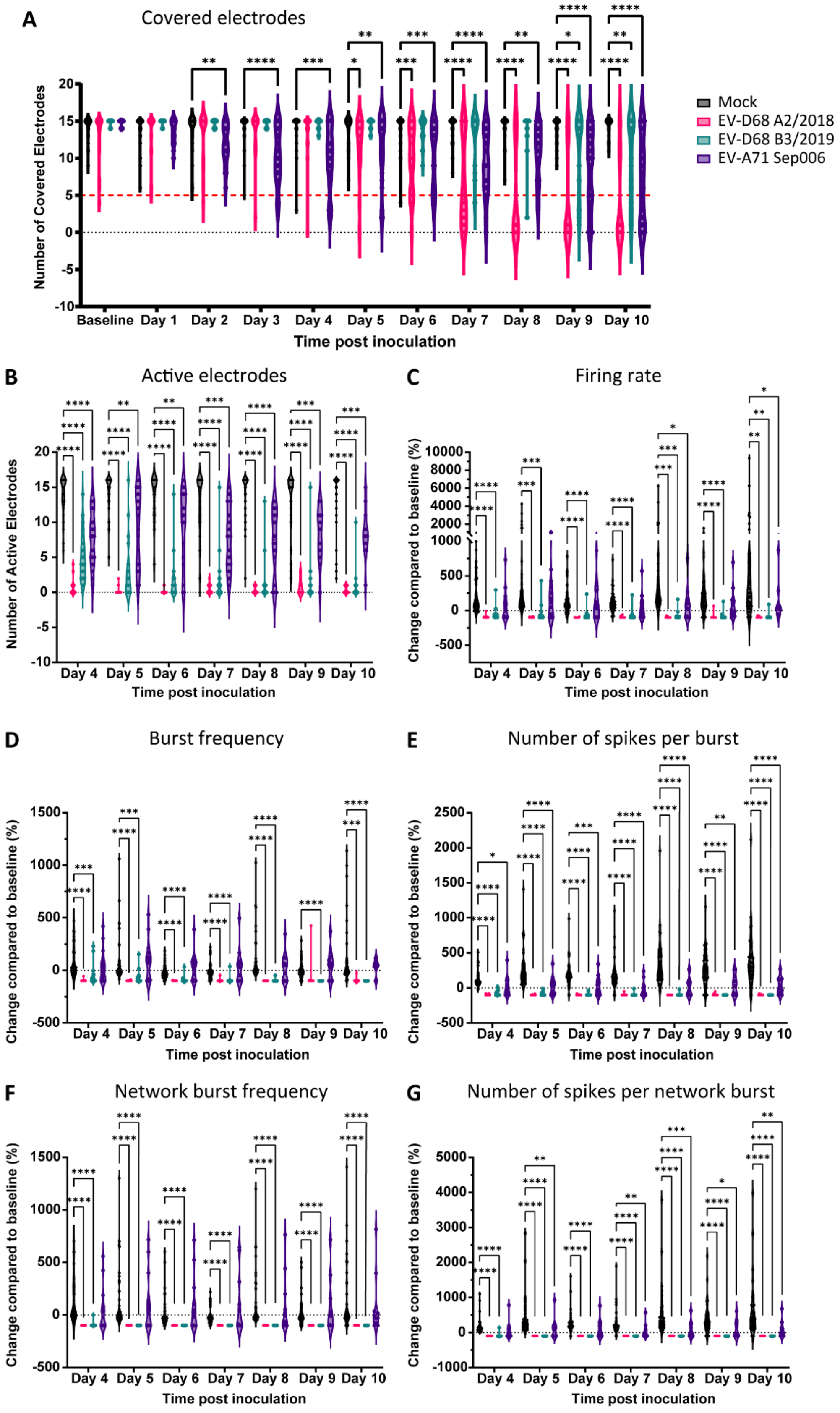

A
Covered electrodes
Number of Covered Electrodes
Baseline Day 1 Day 2 Day 3 Day 4 Day 5 Day 6 Day 7 Day 8 Day 9 Day 10
Time post inoculation
Mock
EV-D68 A2/2018
EV-D68 B3/2019
EV-A71 Sep006
B
Active electrodes
Number of Active Electrodes
Day 4 Day 5 Day 6 Day 7 Day 8 Day 9 Day 10
Time post inoculation
C
Firing rate
Change compared to baseline (%)
Day 4 Day 5 Day 6 Day 7 Day 8 Day 9 Day 10
Time post inoculation
D
Burst frequency
Change compared to baseline (%)
Day 4 Day 5 Day 6 Day 7 Day 8 Day 9 Day 10
Time post inoculation
E
Number of spikes per burst
Change compared to baseline (%)
Day 4 Day 5 Day 6 Day 7 Day 8 Day 9 Day 10
Time post inoculation
F
Network burst frequency
Change compared to baseline (%)
Day 4 Day 5 Day 6 Day 7 Day 8 Day 9 Day 10
Time post inoculation
G
Number of spikes per network burst
Change compared to baseline (%)
Day 4 Day 5 Day 6 Day 7 Day 8 Day 9 Day 10
Time post inoculation

**Supplementary Figure 7. Enterovirus infection impacts the spontaneous activity of co-cultures long-term.** Neural co-cultures were inoculated with EV-D68 A2/2018 (pink), B3/2019 (cyan) or EV-A71 Sep006 (purple) with a MOI of 1, and neural activity was measured between 4 and 10 dpi. The following parameters were displayed (A) number of covered electrodes, where wells were excluded for further analysis if they reached 5 or less covered electrodes throughout the experiment, indicating cell death; (B) number of active electrodes; (C) firing rate; (D) burst frequency, (E) number of spikes per burst, (F) network burst frequency and (G) number of spikes per network burst. Data is displayed from at least four independent experiments performed in six replicates ($n$ = 48 for control, and $n$ = 24 per inoculation group, unless datapoints were excluded based on exclusion criteria, see Material and Methods). Statistical significance was calculated with a two-way ANOVA with a Šídák's multiple comparisons post hoc test. Asterisks indicate statistical significance (*$P$<0.05, **$P$<0.01, ***$P$<0.001, ****$P$<0.0001). Abbreviations: EV = enterovirus; MOI = multiplicity of infection; dpi = days post inoculation.

**References:**


1. Lendemeijer B, Unkel M, Smeenk H, et al. Human Pluripotent Stem Cell-Derived Astrocyte Functionality Compares Favorably with Primary Rat Astrocytes. *eNeuro*. 2024;11(9). doi:10.1523/ENEURO.0148-24.2024
